# Induction of telomerase in p21-positive cells counteracts capillaries rarefaction in aging mice lung

**DOI:** 10.1101/2022.10.27.514005

**Authors:** Larissa Lipskaia, Marielle Breau, Christelle Cayrou, Dmitri Churikov, Laura Braud, Charles Fouillade, Sandra Curras-Alonso, Serge Bauwens, Frederic Jourquin, Frederic Fiore, Rémy Castellano, Emmanuelle Josselin, Carlota Sánchez-Ferrer, Giovanna Giovinazzo, Eric Gilson, Ignacio Flores, Arturo Londono-Vallejo, Serge Adnot, Vincent Géli

**Author notes:** Co-first authors (Larissa Lipskaia, Marielle Breau).

## Abstract

Telomerase is required for long-term cell proliferation and linked to stem cells. This is evident in the lung where short telomeres are associated with lung dysfunction. We constructed a mouse model in which the telomerase (*Tert*) is expressed from the p21^Cdkn1a^ promoter. We found that this peculiar Tert expression curb age-related emphysema and pulmonary perivascular fibrosis in old mice. In old mice lungs, such Tert expression preferentially occurs in endothelial cells where it reduces the number of senescent endothelial cells. Remarkably, we report that Tert counteracts the age-related decline in capillary density. This was associated with an increased number of Cd34+ cells identified as a subclass of capillary cells with proliferative capacity. Expression of catalytically inactive *Tert* neither prevents the decline of capillary density in old mice nor protects against age-related emphysema and fibrosis. These findings reveal that telomerase decreases age-decline of pulmonary functions by sustaining microvasculature regeneration and outgrowth.

## INTRODUCTION

In humans, an increasing number of age-related diseases have been associated with abnormally short telomeres including dyskeratosis congenita, aplastic anaemia, pulmonary fibrosis, and lung emphysema (Armanios, M. & Blackburn, 2012). For never smokers telomere erosion appears as a major risk factor of pulmonary fibrosis, while in smokers mutations that affect telomerase activity favour the development of emphysema and chronic obstructive pulmonary disease (COPD) (Alder *et al*, 2011; Alder *et al*, 2015; Armanios *et al*, 2007; Stanley *et al*, 2014). In mice with critically short or dysfunctional telomeres induction of DNA damage results in the development of pulmonary fibrosis (Povedano *et al*, 2015). In addition, telomerase-deficient mice treated with cigarette smoke develop pulmonary emphysema due to the release of inflammatory cytokines by senescent cells (Alder *et al*, 2011; Amsellem *et al*, 2011; Chen *et al*, 2015). These results inspired development of adeno-associated vectors (AAV) encoding the telomerase reverse transcriptase gene (*TERT*) to assess the therapeutic effect of telomerase ectopic expression either in old mice or in mice with experimentally shortened telomeres (Bernardes de Jesus *et al*, 2012; Piñeiro-Hermida *et al*, 2020; Povedano *et al*, 2018). Telomerase gene therapy had beneficial effects in delaying physiological aging and improving pulmonary function. However, the precise mechanism by which TERT expression prevents lung damage or promotes lung endogenous repair remains to be elucidated.

In response to various stresses including telomere shortening, the p53-dependent up-regulation of p21 expression is thought to be the primary event inducing replicative senescence (Harper *et al*, 1995; Smogorzewska & de Lange, 2002; Herbig *et al*, 2004). p21 is a cyclin-dependent kinase (CDK) inhibitor that interacts with multiple CDK complexes, with a higher affinity for CDK4/2, involved in the G1/S transition (Harper *et al*, 1995). In normal conditions, the levels of p21 are also regulated by several additional transcription factors including E2F1, STAT3, and MYC (Abbas & Dutta, 2009). One major effect of p21 in stem cell compartments is to halt cell proliferation. Reciprocally, mice with p21 deletion present an increased number of stem cells under normal homeostatic conditions, however their number declines with age suggesting that p21 is important for their life-long maintenance (Cheng *et al*, 2000; Kippin *et al*, 2005). Since dysfunctional telomeres signal cell cycle arrest via the ATM-p53-p21 pathway (Smogorzewska & de Lange, 2002; Herbig *et al*, 2004), we generated a transgenic model in which the telomerase is expressed under the p21 promoter regulation. The p21^+/Tert^ model is expected to up-regulate *Tert* expression in response to telomere dysfunction, but also in response to other cues inducing p21. Age-related decline of pulmonary function is a paradigm for telomere dysfunction-driven degenerative process characterized by p21 and p16 activation and cellular senescence (Barnes *et al*, 2019). We uncovered that p21 promoter-dependent expression of TERT does promote lung function in aged mice, and this coincides with maintenance of microvascular density, decreased senescence of endothelial cells, and preservation of endothelial cells endowed with proliferative capacities in the lungs of old mice.

## RESULTS

### Generation and validation of the p21^+/Tert^ mouse model

To generate a mouse model that expresses TERT under control of the p21^Cdkn1a^ promoter, we substituted the start codon of the *Cdkn1a* gene by a *mCherry-2A-Tert* cassette (Fig. 1A). The detailed construction of the targeting vector and integration of the cassette are shown in the *figure source data*. The *mCherry-2A-Tert* allele retains regulatory 5’ and 3’UTRs of the endogenous *Cdkn1a* gene. Translation of the polycistronic *mCherry-2A-Tert* mRNA produces two separate mCherry-2A and mTert polypeptides due to the ribosome skipping at the 2A sequence (Fig. 1A). The generated p21^+/Tert^ mouse therefore produces p21 protein from one allele and mCherry and TERT from the other. We created in addition another mouse line (p21^+/Tert-CI^) identically designed but with a point mutation (D702A) in the active site of TERT which abolishes telomere elongation by telomerase (Barnes *et al*, 2019; Lingner *et al*, 1997) (Fig. 1A). The functionality of TERT expressed from the *mCherry-2A-Tert* cassette was validated *in vitro* by checking its ability to elongate telomeres in mouse ES *Tert-/-* cells (Fig. 1B). p21 induction was further verified in p21^+/Tert^ mice by following the whole body fluorescence emitted by the mCherry after exposure to ionising radiation, a condition known to induce p53-dependent expression of p21 (Fig. EV1A). A similar validation was performed after the treatment of p21^+/Tert^ mice with doxorubicin by following the expression of the cassette in liver and kidney that both concentrate doxorubicin (Fig. EV1B).

**Figure 1.**
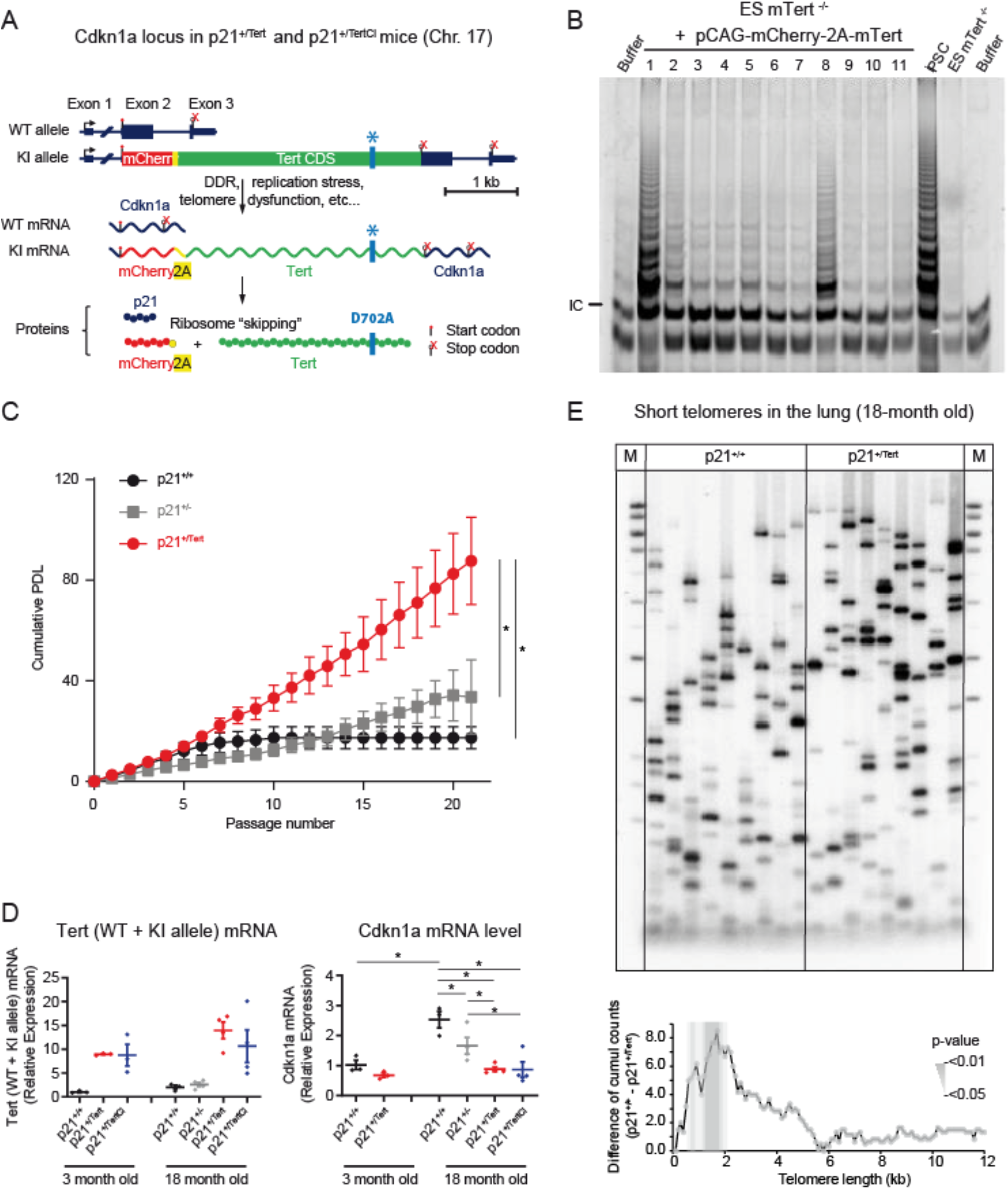
Construction and validation of the p21^+/Tert^ mouse model. **(A)** Schematic of the modified *Cdkn1a* locus, the mRNAs transcribed, and the proteins translated. The *mCherry-2A-Tert* cassette was inserted in place of the start codon of Cdkn1a (see source data for details). The gene locus is drawn to scale with intron 1 contracted. The 2A peptide sequence causes a “ribosomal skipping” that generates two independent polypeptides, mCherry and TERT, from the same mRNA. In the *mCherry-2A-Tert*^*CI*^ cassette the codon GAT encoding D702 essential for Tert catalytic activity has been replaced with GCA encoding A702. **(B)** To demonstrate that TERT expressed from the *mCherry-2A-Tert* cassette is functional, we transfected Tert^−/-^ ES cells with a plasmid carrying *mCherry-2A-Tert* under control of the constitutively active pCAG promoter and checked telomerase activity *in vitro. In vitro* telomerase activity was assayed using Telomere Repeat Amplification Protocol (TRAP). The 6 bp ladder reflects telomerase activity. The arrow indicates the PCR internal control (IC). iPS and ES Tert^−/-^ cells were used as positive and negative controls, respectively. **(C)** p21-promoter dependent TERT expression bypasses senescence in PA-SMCs *ex-vivo*. Cumulative population doubling level (PDL) of PA-SMCs isolated from mice from the three mentioned genotypes. The data points are the means of 8 independent cultures established from individual mice. *p<0,05 comparing p21^+/Tert^ with p21^+/+^ and p21^+/-^, student t-test. **(D)** Left panel, relative expression of Tert mRNA (endogenous + transgene) in the whole lung samples measured by RT-qPCR. *Tert* expression is shown separately for young and old mice. Right panel, relative expression of *Cdkn1a* (p21) measured by RT-qPCR in the same lung samples. **(E)** Telomere Shortest Length Assay (TeSLA) performed on lung parenchyma from p21^+/+^ and p21^+/Tert^ mice. The left panel depicts 2 representative Southern blots probed for the TTAGGG repeats while the right panel shows the difference of cumulative number of short telomeres between lungs from p21^+/+^ and p21^+/Tert^ mice. This difference is significant for telomeres size range between 0.4 and 2.0 kb. Southern blots for all mice are shown in Fig. EV3.

To further validate p21-promoter dependent expression and function of TERT during replicative senescence, we examined the proliferation of cultured lung pulmonary artery smooth muscle cells (PA-SMCs) isolated from young wild type and transgenic mice (Fig. 1C). Pulmonary artery smooth muscle cells (PA-SMCs can be readily isolated from mouse lungs and cultured. As many other mouse cell types, PA-SMCs senesce quickly in standard cell culture conditions (21% atmospheric oxygen) and cell proliferation arrest is thought to be caused by oxidative DNA damage (Parrinello *et al*, 2003), possibly by altering the very long mouse telomeres. We found that PA-SMCs isolated from p21^+/+^ and p21^+/-^ mice entered senescence after a few passages, with a final mean population doubling levels (PDLs) of 15,51 (+/-4,14) and 36,09 (+/-8,30) respectively, the difference between the two mice being likely due to p21 haplo insufficiency. In contrast, the cells from p21^+/Tert^ mice never entered senescence and proliferated at a much faster rate (Fig. 1C). The difference in cumulative PDL between p21^+/Tert^ and both p21^+/+^ and p21^+/-^ became significant at passage 7. We confirmed the expression of the mCherry-2A-Tert transcript by RT-qPCR (Fig. EV2A, B), and also detected a slight increase of telomerase activity at passages 2-4, before PA-SMCs cumulative PDL curve became significantly different (Fig. EV2C). We measured the load of very short telomeres (VSTs) that may arise in cultured PA-SMCs causing their arrest by TeSLA (Lai *et al*, 2017) (Fig. EV2D). While the cumulative number of VSTs increased with passages in p21^+/+^ cells, it did not change in p21^+/Tert^ cells (Fig. EV2E). Thus, we concluded that mouse cultured mouse cells do accumulate VSTs coincidently with proliferation arrest, and that ectopic Tert expression can effectively counteract accumulation of VSTs coincidently with abrogation of the arrest. We next analysed Tert (and Tert^CI^) as well as p21 expression in the lungs of 18 month-old mice (Fig. 1D). We found that Tert and Tert^CI^ were expressed in p21^+/Tert^ and p21^+/TertCI^ mice, respectively (Fig. 1D, left panel). Interestingly, p21 transcript was reduced in the lungs of old p21^+/Tert^ mice relative to age-matched p21^+/+^ controls (Fig. 1D, right panel). To our surprise, p21 transcript level was also reduced in the lungs of the p21^+/TertCI^ suggesting that p21 expression is attenuated in a way that is independent of its catalytic activity (see the discussion).

We finally asked whether Tert controlled by p21 promoter would also curb the accumulation of VSTs in the lung parenchyma of old mice by measuring the load of VSTs in the lung parenchyma of 18 month-old p21^+/+^ and p21^+/Tert^ mice (Fig. 1E and Fig. EV3). We found that lungs from p21^+/Tert^ compared to p21^+/+^ mice harboured significantly less VSTs in the 0.4 -2.0 kb range (Fig. 1E) suggesting that p21-promoter dependent Tert expression from in old mice is able to partially heal critically short telomeres *in vivo*. Overall, p21-promoter expression of Tert is associated with a decrease of VSTs, however and surprisingly reduction of p21 levels is separable from telomere elongation since it occurs in p21^+/TertCI^ mice.

### p21 promoter-dependent expression of Tert protects against age-related emphysema and perivascular fibrosis

We sought to determine the impact of the conditional expression of Tert on age-related manifestations in the lung, such as emphysema and fibrosis. C57BL/6NR mice used in this study naturally develop age-related emphysema and mild fibrosis. Emphysema is characterized by lung airspace enlargement as a consequence of a decrease of lung elasticity with advanced age (Sharma & Goodwin, 2006). Morphometry studies (Dunnill, 1962) did not reveal air space enlargement in young mice (Fig. 2A, B). In contrast, old p21^+/+^, p21^+/-^, and p21^+/TertCI^ mice developed emphysema reflected by an increase of alveolar size (measured by the mean-linear intercept or MLI). Remarkably, old p21^+/Tert^ mice did not exhibit air space enlargement (Fig. 2A, 2B). The fact that p21^+/Tert-CI^ mice exhibit the same aging characteristics in the lung as control mice supports the idea that TERT’s effect in emphysema protection is related to its ability to elongate critically short or damaged telomeres (Matmati *et al*, 2020; Birch *et al*, 2015; Fouquerel *et al*, 2019).

**Figure 2.**
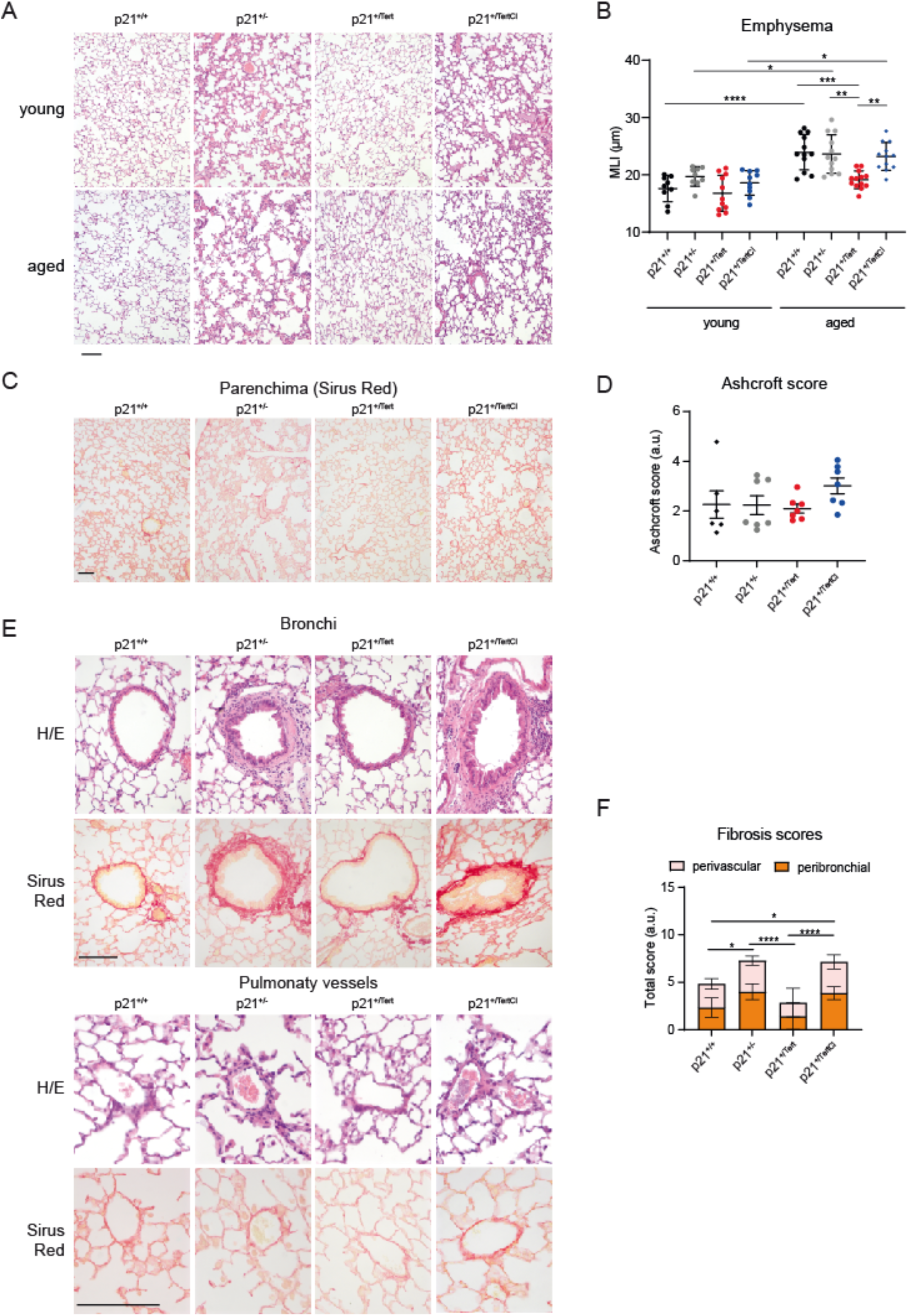
Telomerase protects against age-related emphysema and perivascular fibrosis. **(A)** (Representative micrographs showing lung parenchyma of young and old mice from the four mouse models stained with hematoxylin/eosin. **(B)** Scatter plot showing mean liner intercept (MLI) measurements in young and old mice. High MLI values indicate development of emphysema-like phenotype. **(C)** Representative micrographs showing lung parenchyma of aged mice of the 4 genotypes stained with H/E or Sirus Red used to visualize collagen deposition (hallmark of lung fibrosis). **(D)** Scatter-plot graph showing parenchymal fibrosis quantification according to Aschcroft score. (**e**) Representative micrographs showing bronchi and pulmonary vessels of aged mice of the 4 genotypes stained with H/E or Sirus Red. (**F**) Bar-graph showing perivascular and peribronchial fibrosis scoring. Fibrosis scores were attributed on 1 - 5 scale: 0-absent; 1-isolated mild fibrotic changes, 2-clearly fibrotic changes; 3-substantial fibrotic changes, 4 – advanced fibrotic changes, 5-confluent fibrotic masses. For all graphs, *P<0.05, **P<0.01, ***P<0.001, ****P<0.0001 (One way ANOVA with Bonferroni correction). For all images, Bar - 50 μm.

We also searched for manifestations related to lung fibrosis in the mouse models. We did not observe any differences in fibrosis in the parenchyma among the four mouse models (Fig. 2C, 2D). However, Sirus Red and HE stainings revealed local increase of fibrosis around bronchi and vessels in lungs from the old p21^+/-^ and p21^+/TertCI^ mice compared to WT (Fig. 2E and 2F). In contrast, fibrosis was decreased in lungs from p21^+/Tert^ mice compared to WT (Fig. 2E and 2F).

Collectively, these results indicate that p21-dependent expression of TERT protects mice from agerelated emphysema and locally from perivascular and bronchial fibrosis.

### p21 is preferentially expressed in lung endothelial cells

To understand the effects associated with the p21-promoter dependent expression of *Tert* in lung, we determined in which lung cell types p21 is expressed. To this end, we performed single-cell RNA sequencing (scRNA-seq) on whole lungs from 18 month-old p21^+/+^, p21^+/-^, p21^+/Tert^ and p21^+/TertCI^ mice. We performed dimensionality reduction and unsupervised cell clustering to identify distinct cell types based on shared and unique patterns of gene expression (see Methods). For each mouse, clustering of gene expression matrices identified cell types that were in good agreement with the published Mouse Cell Atlas (MCA) (Han *et al*, 2018) (Fig. 3A). As shown in Fig. EV4 and Table 1, similar cell types were isolated from the lung of the mice of the four genotypes (WT (n=1); p21^+/-^ (n=3), p21^+/Tert^ (n=5) and p21^+/TertCI^ (n=2)) suggesting that Telomerase expression does not change drastically the cell population composition in 18 mo mice.

**Figure 3.**
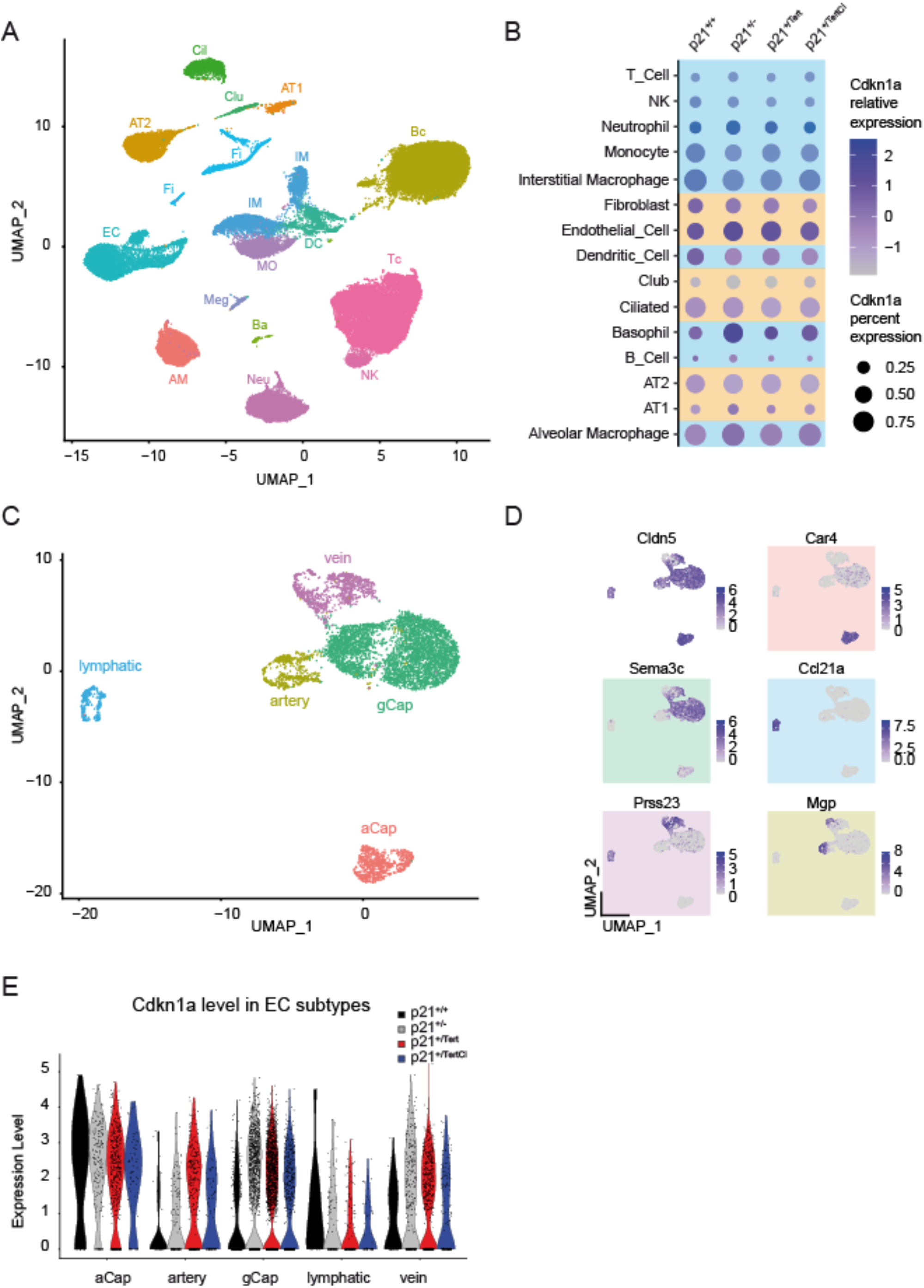
p21 is preferentially expressed in lung endothelial cells. **(A)** Unsupervised Uniform Manifold Approximation and Projection for Dimension Reduction (UMAP) clustering of lung cells. Lung cell populations were identified in lung samples from WT (p21^+/+^), p21^+/-^, p21^+/Tert^ and p21^+/TertCI^ 18 month-old mice using Mouse Cell Atlas (MCA) annotation procedure. **(B)** Dot plots of Cdkn1a expression in the different lung cell-types. The identified cell types are shown on the y-axis. The size of the dots represents the fraction of the cells expressing Cdkn1a. The color intensity represents the average expression level in p21-positive cells. Immune and constitutive lung cells are color-coded in blue and orange, respectively. **(C)** UMAP clustering of lung endothelial cells from WT (p21^+/+^), p21^+/-^, p21^+/Tert^ and p21^+/TertCI^ 18 month-old mice. Cell populations were identified based on known markers for these endothelial cell subtypes^26^. (Art) artery EC cells; (Cap_1; also called aCap) capillary 1 EC cells; (Cap_2; also called gCap) capillary 2 EC cells^27^; (Lym) lymphatic EC cells and (Vein) vein EC cells. **(D)** Representative markers used to annotate lung endothelial cell classes. **(E)** Violin plots showing p21 expression. ECs are subdivided into groups according to cluster and genotype (WT (black); p21^+/+^ (grey), p21^+/Tert^ (red) and p21^+/Tert CI^ (red). For all panels WT (n=1), p21^+/-^ (n=3), p21^+/Tert^ (n=5), p21^+/TertCI^ (n=2).

We then analyzed p21 expression in different cell types in mice of the four genotypes. Among immune cells, p21 was mainly expressed in interstitial and alveolar macrophages, monocytes, and in dendritic cells, while among non-immune lung cells, it was mainly expressed in endothelial cells (ECs) as compared to other cell types including AT1 and AT2 cells (Fig. 3B). Unfortunately, we could not directly detect the transgene (mCherry-2A and Tert) transcript because the 10x Genomics single-cell 3′ chemistry only provides sequence information on the short region preceding the polyA-tail. Since p21 (and hence Tert and Tert^CI^ in p21^+/Tert^ and p21^+/TertCI^ mice, respectively) is preferentially expressed in pulmonary ECs we focused our single-cell analysis on this cell type. We annotated lung ECs to the specific vascular compartments based on recently published markers of lung ECs (Kalucka *et al*, 2020). We identified 5 classes of EC comprising capillary, artery, vein, and lymphatic classes (Fig. 3C, D). Capillary class 1 was recently identified as aerocytes (aCap) involved in gas exchange and trafficking of leucocytes, and capillary class 2 as general capillary cells (gCap) that function in vasomotor tone regulation and in capillary homeostasis (Gillich *et al*, 2020). We uncovered that p21 is expressed at the highest level in aerocytes (aCap) (Fig. 3E). This result is in agreement with Tabula Muris Senis (https://tabula-muris.ds.czbiohub.org) (Fig. EV5). In aCap, we also observed that p21 expression was reduced in both p21^+/Tert^ and p21^+/TertCI^ mice but this was not observed in the other endothelial cell sub-types (Fig. 3E).

### Tert expression counteracts lung senescence of endothelial cells

We next investigated whether TERT could suppress age-related cellular senescence in lungs. For this, we analysed the colocalization of the conventional senescence marker p16 with either the endothelial marker CD31 (Pecam-1) or the alveolar epithelial type II cell marker Muc1 in lung sections from 18-mo-old mice of the 4 genotypes (Fig. 4). Single-cell RNA-seq carried out on lungs of the 4 mouse models confirmed the specificity of these markers (Fig. EV6). Importantly, we validated anti-p16 antibody by checking that p16 staining detected after an injury (leading to the induction of p16) disappeared after inducible elimination of the cells expressing p16 (see Born *et al*, 2022). We found that only 10 % of the p16-positive cells expressed Muc1 and p21-dependent expression of TERT did not reduce the fraction of p16-positive cells marked by Muc1 (Fig. 4A and 4B). In contrast, we found that about 30% of the p16-positive cells were also positive for the endothelial marker CD31 (Fig. 4C, see the higher magnification and 4D) indicating that endothelial cells largely contributed to senescence in the lungs of the old mice. Remarkably, p21^+/Tert^ lung samples accumulated significantly less p16-positive CD31 cells (Fig. 4C and 4D) suggesting that TERT expression counteracts senescence of endothelial cells or that senescent endothelial cells are more efficiently eliminated in these mice. To further confirm the senescence suppression by the conditional expression of TERT, we also assessed the global accumulation of senescent cells in the lungs of young and 18-month-old mice of the 4 genotypes by evaluating the number of lung parenchymal cells with senescence associated β-galactosidase activity (SA-β-Gal) (Fig. EV7A, 7B). While WT (p21^+/+^), p21^+/-^, and p21^+/TertCI^ mice displayed a significant increase in the percentage of SA-β-Gal with age, the percentage of SA-β-Gal-positive cells increased only slightly in p21^+/Tert^ mice.

**Figure 4.**
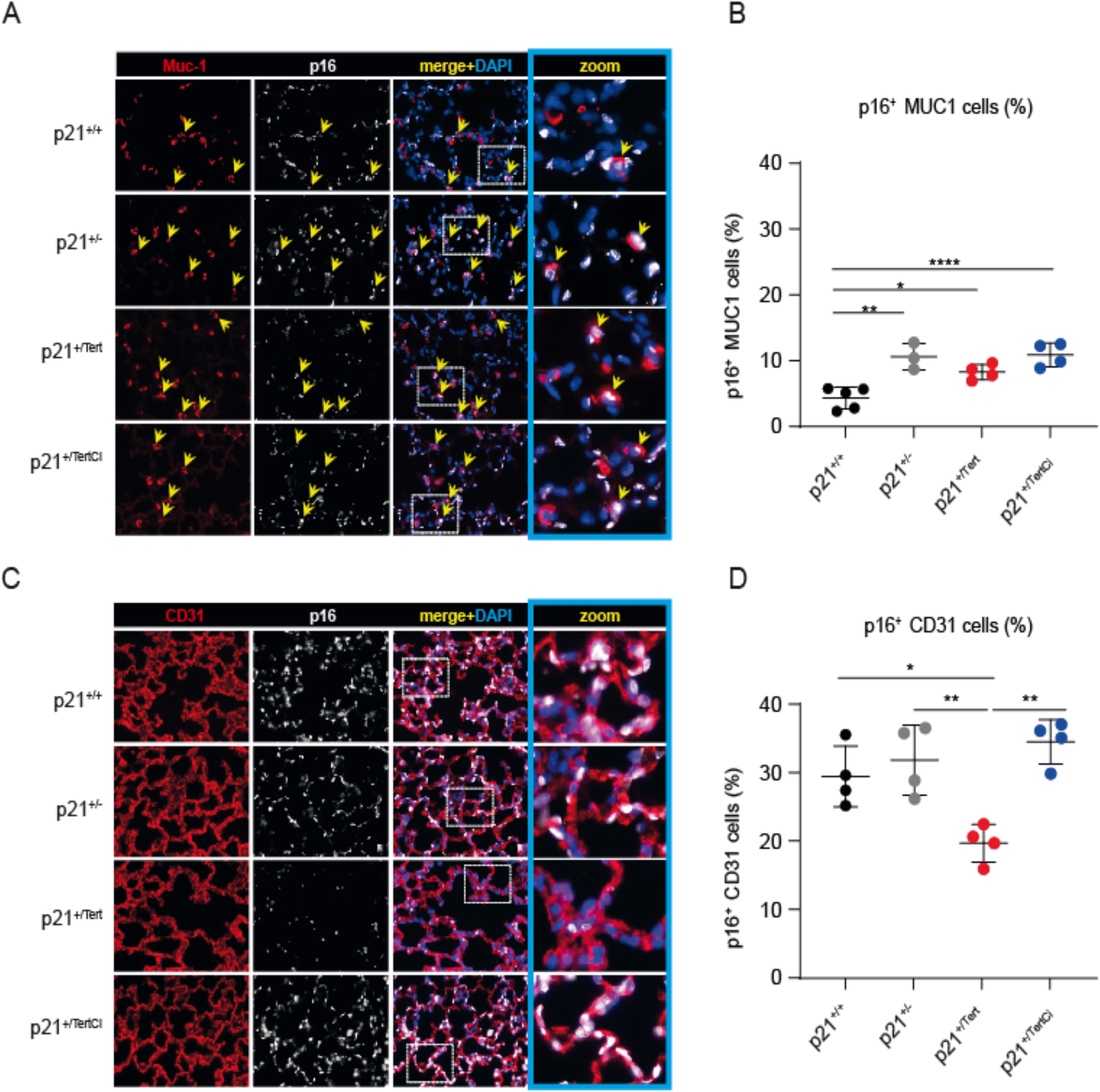
Endothelial cells senescence is attenuated in aged p21^+/Tert^ mice. **(A)** Representative micrographs showing immunofluorescence of p16 (white) in lung cells co-stained either with Muc1 (red, a marker of AT2 cells). Arrows indicate cells co-stained for both Muc1 and p16. The zoomed areas are indicated. Blue rectangle – DAPI nuclear staining. Bar - 50 μm. **(B)** Scatter-plot graphs representing the percentage of p16^+^ Muc1^+^ cells in the different groups of mice. *P<0.05, **P<0.01, ***P<0.001. (One way ANOVA with Bonferroni post hoc test). **(C)** Same as (**a**) except that p16 lung cells (white) are co-stained with Cd31 (red, a marker of endothelial cells, lower panel). **(D)** Same as (**b**) except that the graph represents the percentage of p16^+^ Cd31+ cells.

### TERT expression counteracts the age-related decline in capillary density and maintains a high number of CD34+ cells in the lungs of old mice

Microvasculature density has been shown to decrease in organs of aged mice (Grunewald *et al*, 2021). Because conditional induction of TERT has been reported in vivo to cause the long-term proliferation of several types of stem cells (Sarin *et al*, 2005; Shkreli *et al*, 2011; Montandon *et al*, 2022), we determined whether TERT expressed from p21 promoter in the lungs of aged mice could affect capillary density that declines with age. We labelled lung parenchyma with CD31 to reveal the vasculature compartment in young and old mice of the 4 genotypes (Fig. 5A). Quantification of the labelling revealed a higher microvasculature density in the aged p21^+/Tert^ mice compared to the 3 other genotypes (Fig. 5B). These results demonstrate that Tert (but not Tert^CI^) counteracts the decline in capillary density in the lungs of aged mice.

**Figure 5.**
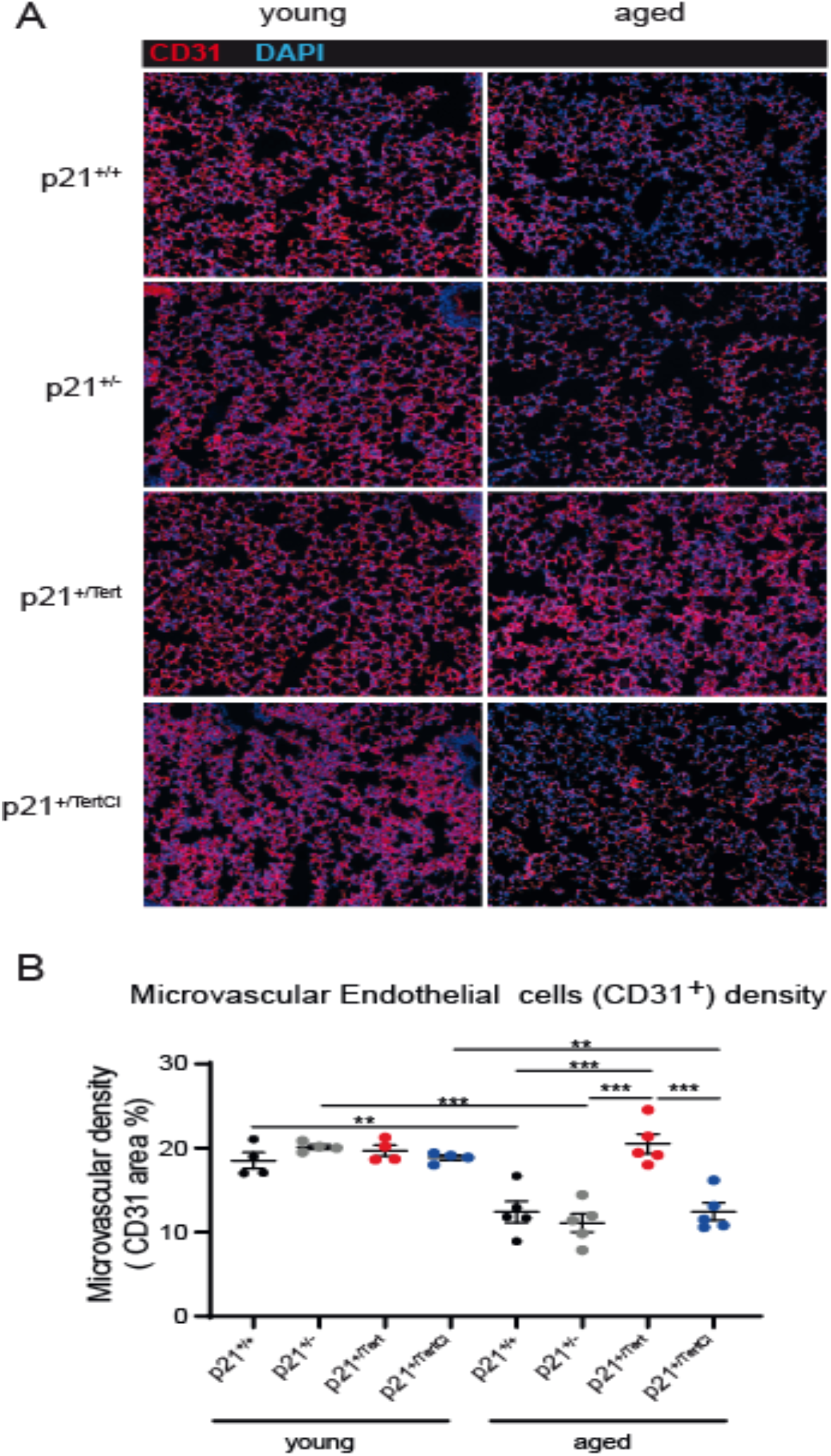
TERT expression in p21-positive cells counteracts age-related decline in capillary density. **(A)** Representative micrographs showing immunofluorescence of CD31 (red, a marker of endothelial cells). Bleu – DAPI nuclear staining. Bar - 50 μm. **(D)** Scatter-plot graph showing microvascular density in different groups of young (4 m) and old (18 m) mice. *P<0.05, **P<0.01, ***P<0.001, ****P<0.0001 (One way ANOVA with Bonferroni post hoc test).

We next thought to label in the lung of old mice CD34 that was found to mark a subtype of capillary cells involved in healing capillaries after acute lung injury (Ding *et al*, 2011; Niethamer *et al*, 2020; Wang *et al*, 2022). We thus examined lung parenchyma from young and old mice for the presence of cells marked by CD34. Staining of lung sections from young and old mice for the presence of cells expressing CD34 revealed that all young mice harboured more CD34+ cells as compared to the old mice (Fig. 6A). However, with age, CD34^+^cells were hardly detected except for the p21^+/Tert^ mice in which the level of CD34^+^ cells was similar to the level observed in young mice (Fig. 6B). At lower magnification, we could clearly observe a population of Cd34+ cells surrounding vessels in old WT and p21^+/Tert^ mice and another population of Cd34+ in the lung parenchyma mainly present in the old p21^+/Tert^ mice (Fig. EV8).

**Figure 6.**
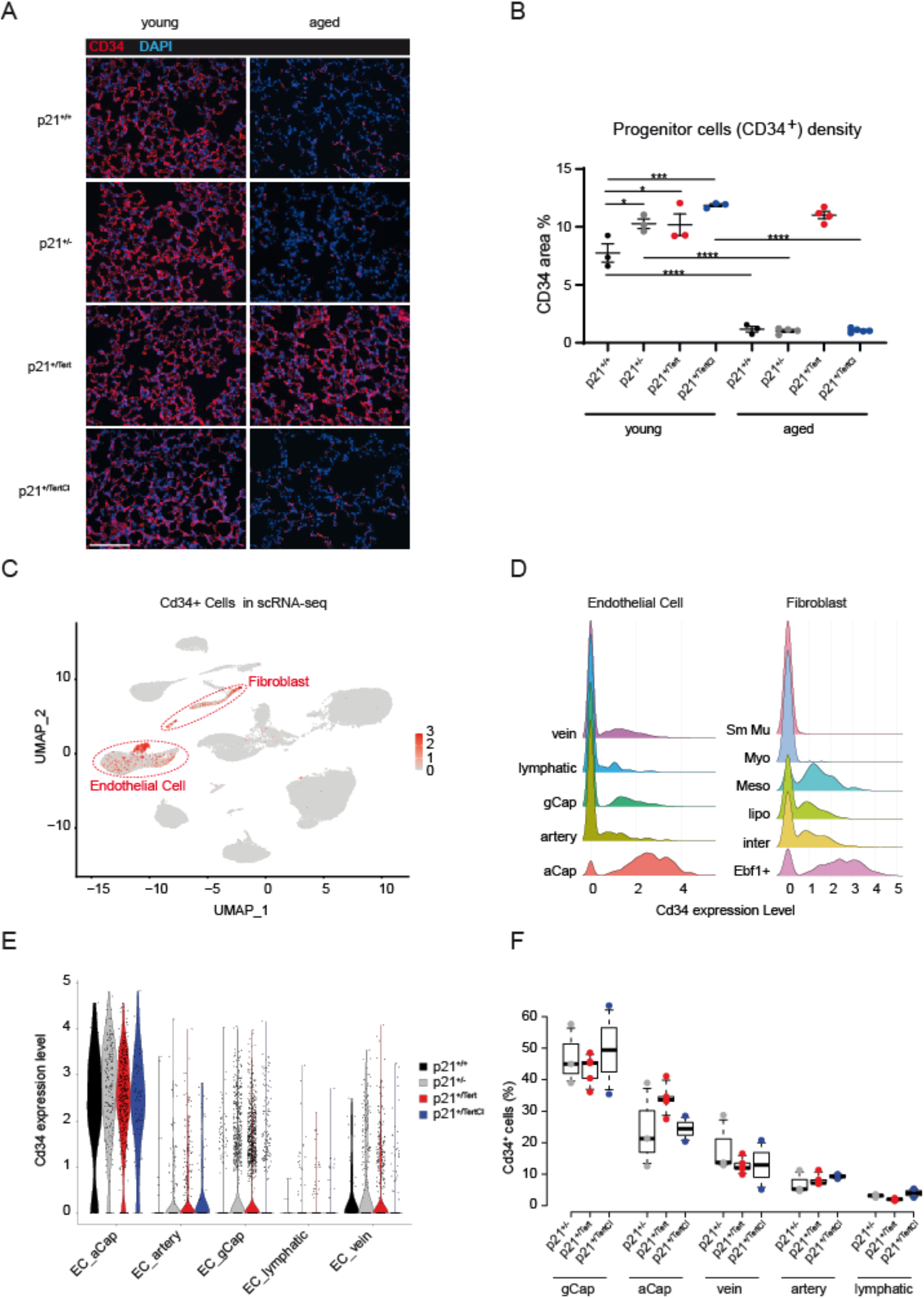
TERT expression maintains a high number of CD34+ cells in the lungs of old mice. **(A)** Representative micrographs showing immunofluorescence of CD34 (red). DAPI nuclear staining (blue). Bar - 50 μm. **(B)** Scatter-plot graph showing lung area of CD34 expression in different groups of young (4 m) and old (18 m) mice. *P<0.05, **P<0.01, ***P<0.001, ****P<0.0001 (One way ANOVA with Bonferroni post hoc test). **(C)** UMAP clustering of lung cells from 1xWT, 3xp21^+/-^, 5xp21^+/Tert^ and 2xp21^+/TertCI^ mice. Cells expressing Cd34 gene are visualised by red dots (negative cells are in grey). **(D)** Ridget plot of Cd34 expression level for each subtype of EC and Fibroblast cell types. **(E)** Violin plots representing the expression (log(counts)) of Cd34 in EC subtypes for each genotype **(F)** Boxplot of the distribution of Cd34+ cells in each EC subtype relative to the total number of Cd34+ ECs. Individual mice are represented. p21^+/-^(x3, grey), p21^+/Tert^ (x5, red) and p21^+/TertCI^ (x2, blue).

Overall, old p21^+/Tert^ mice maintained this high level of CD34-expressing cells while it dropped drastically in aged mice of all other genotypes. These results raised the question of the identity of the CD34+ cells in the lungs from 18-month old mice.

We turned to scRNA-seq of the lungs from 18-month-old p21^+/+^, p21^+/-^, p21^+/Tert,^ and p21^+/TertCI^ mice. We found that Cd34 was mainly expressed in EC (64.9% of Cd34+ lung cells) and in fibroblasts (26.2% of Cd34+ lung cells) (Fig. 6C). We further look at the expression of Cd34 in the different subclasses of EC and fibroblasts. We found that Cd34 was expressed at highest levels in Ebf1+ fibroblasts and mesothelial cells (>85% are Cd34+ cells), two populations of fibroblasts delineated by early B-cell factor 1 and Upk3b expression, respectively (Sidney *et al*, 2014; Korsunsky *et al*, 2022) (Fig. 6D). In EC, Cd34 expression was maximum in aCap (aerocytes; >90% are Cd34+ cells) but also expressed in gCap and vein cells but at lower levels (between 25% to 29% are Cd34+ cells respectively) (Fig. 6D). However, mRNA level of Cd34 in the different classes of lung ECs of the 4 genotypes did not reveal major differences (Fig. 6E), in apparent contradiction with the immunostaining (see discussion). We thus sought to compare the distribution of Cd34+ positive cells in p21^+/-^, p21^+/Tert^, and p21^+/TertCI^ mice (Fig. 6F). We found that in comparison with p21^+/-^ and p21^+/TertCI^ mice, p21^+/Tert^ mice have a higher number of aCap cells expressing Cd34 (Fig. 6F and Fig. EV9).

### CD34+ endothelial cells show proliferative capacity in the lungs of old mice

We sough to determine whether the CD34+ cells were endowed with proliferative capacity. We stained lung cells with antibodies directed with CD34 and PCNA. We found that p21^+/Tert^ old mice had a higher number of CD34+ cells labeled with the proliferation marker PCNA compared to the control mice while this difference was much less pronounced in young mice (Fig. 7A, left and right panels). To further assess CD34+ cell proliferation more directly, 18-month-old mice were injected with Bromodeoxyuridin (BrdU) *intra*-*peritoneally* 24h before lung sampling. Although the number of BrdU positive cells was low, we found a higher number of BrdU positive cells in the lungs of p21^+/Tert^ mice compared to the WT (p21^+/+^), p21^+/-^, and p21^+/TertCI^ mice (Fig. 7B, C). Interestingly, most of the BrdU-positive cells were also labelled with CD34 (Fig. 7C). These results show that p21 promoter-dependent expression of TERT endows CD34+ cells with proliferative capacity.

**Figure 7.**
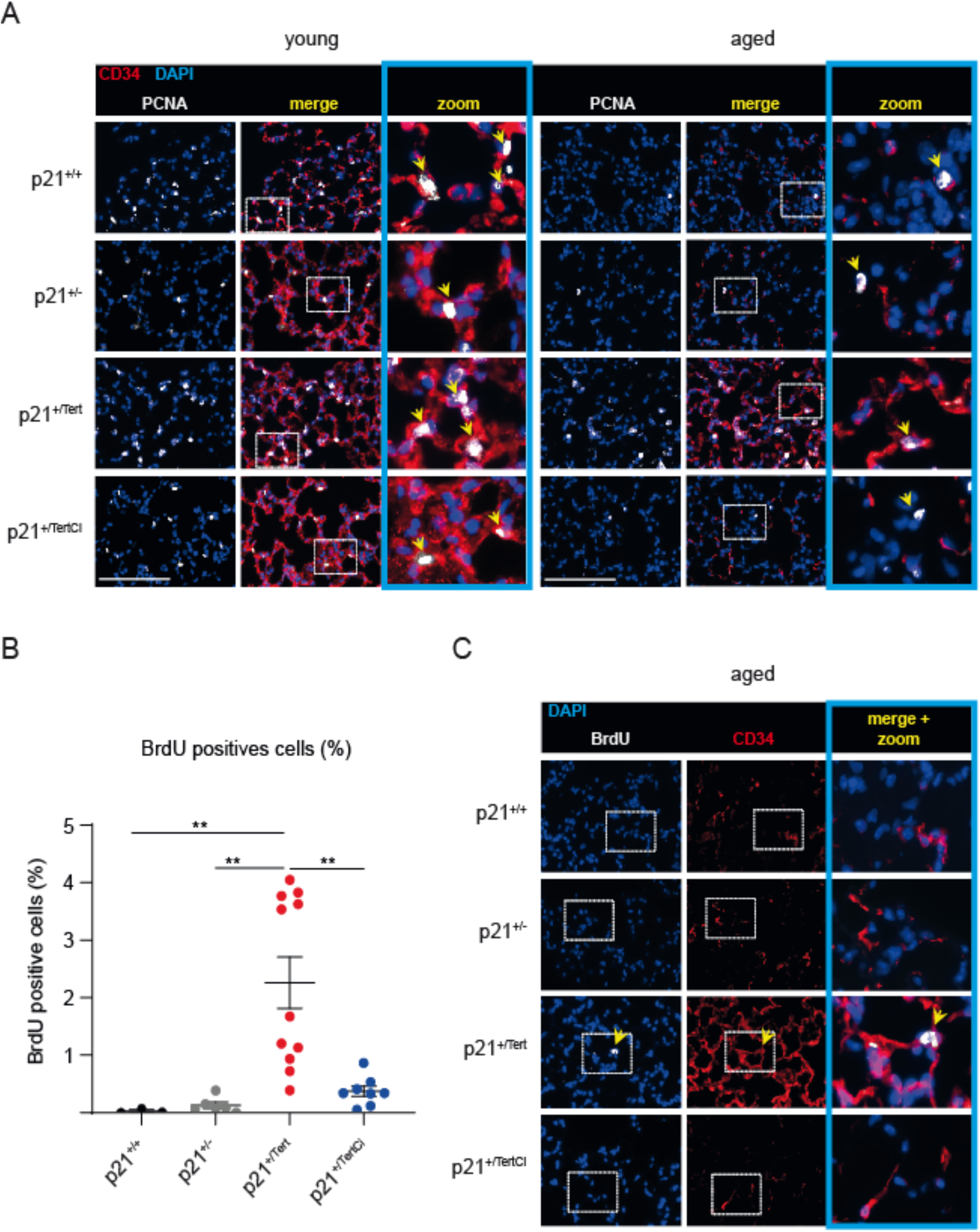
Lung CD34^+^ cells preserve their capacity to proliferate in aged p21^+/Tert^ mice. (**A**). Representative micrographs of mice lung showing proliferating cells (arrows in zoomed areas) identified by nuclear immunofluorescence of PCNA (white, proliferating cells nuclear antigen) in lung of young and old mice of all genotypes. The majority of proliferating cells were also co-stained with CD34 (red, a marker of endothelial cell precursor). Blue – DAPI nuclear staining. The zoomed areas are indicated. Bar - 50 μm. **(B)** Scatter-plot graph showing percentage of BrdU-positive cells in different types of mice. **P<0.01. (One way ANOVA with Bonferroni post hoc test). **(C)** 18-mo-old mice were injected with BrdU intra-peritoneally 24h before organ sampling. Representative micrographs of mice lung showing CD34^+^ cells (red) co-stained with anti-BrdU antibody (white). Bleu – DAPI nuclear staining. Arrows indicate CD34^+^ cells incorporating BrdU. The zoomed areas are not indicated. Bar - 50 μm.

## DISCUSSION

Numerous studies have shown that abnormally short telomeres and senescent cell accumulation are found in the lungs of COPD patients suggesting a direct relationship between short telomeres, senescence and the development of the disease (Alder *et al*, 2011; Stanley *et al*, 2014; Houben *et al*, 2009; Savale *et al*, 2009; Tsuji *et al*, 2010). These conclusions have been strengthened by mouse studies showing that alveolar stem cell failure is a driver of telomere-mediated lung disease (Alder *et al*, 2015). Collectively these results suggest that DNA damage, cellular senescence, stem cell exhaustion, and likely mitochondrial dysfunction, all induced by accumulation of short telomeres, contribute together to lung functional demise. Because TERT may reverse these processes, TERT-based gene therapy (Martínez & Blasco, 2017) may be clinically beneficial to treat COPD. However, the mechanism by which *in vivo Tert* expression could prevent the occurrence of emphysema during aging remained to be elucidated.

In this study, we designed a fine-tuned regulatory loop allowing the expression of telomerase under conditions that induce the expression of p21, including the accumulation of critically short telomeres. Importantly this conditional expression of TERT has been introduced in telomerase-positive mice (Tert^+/+^), e.g., with long telomeres underscoring the importance of few critically short telomeres even in WT mice. Although p21 expression is not only dependent of p53, the p21^+/Tert^ mouse provides a unique tool allowing us to evaluate the impact of TERT re-expression on age-related diseases. We exploited the fact that C57BL/6NR mice naturally develop age-related emphysema and mild fibrosis to test *in vivo* the specific effects of TERT expression.

Our results indicate that catalytically active TERT is able to sustain proliferation of cells marked with CD34+ cells throughout life, and this correlates with the maintenance of capillary density during aging. Remarkably, the p21 promoter-driven expression of TERT has several related consequences in aging lungs: 1) reduction of the age-dependent accumulation of senescent cells, many of them being endothelial cells, 2) promoting cell proliferation, particularly of the CD34+ endothelial cells 3) life-long maintenance of the lung capillary density, and 4) attenuation of age-related emphysema and perivascular fibrosis. Strikingly, we never observed lung tumors in the p21^+/Tert^ mice.

Collectively, these results lead us to propose that maintenance of capillary density in aged mice linked to the sustained proliferation of CD34+ cells could protect lungs against age-related emphysema and perivascular fibrosis. The fact that TERT could prevent age-related emphysema by promoting micro-vasculature is supported by previous studies showing a tight relationship between vascular density, endothelial cells function and emphysema (Grunewald *et al*, 2021; Cordasco *et al*, 1968; Kasahara *et al*, 2000; Kasahara *et al*, 2001; Rafii *et al*, 2021). It was recently shown that VEGF signalling was greatly reduced in multiple organs during aging, and that supplying VEGF protected age-related capillary loss^34^. Whether increased VEGF levels in the lungs of the aged p21^+/Tert^ mice contribute to the protective effects of the conditional expression of TERT remains to be determined.

We found that inactivation of telomerase catalytic activity abolishes the protection conferred by the transgene, thus pointing out the crucial role of TERT canonical activity in sustaining lung function during aging. This is in agreement with studies indicating that short telomeres are important for the predisposition to lung diseases in patients with mutations affecting telomere elongation by telomerase (Stanley *et al*, 2014). Interestingly, we detected a correlation between the presence of very short telomeres (<400 bp) and cellular senescence in the lung. We thus assume that induction of the Tert expression from activated p21 promoter might repair damaged telomeres. Indeed, it has been previously reported that the DNA damage response at telomeres was contributing to lung aging and COPD (Birch *et al*, 2015). However, we found that expression of both Tert and Tert^CI^ globally decrease the levels of p21 in the lungs. This is not due simply to the haploinsufficiency of p21 in the p21^+/Tert^ and p21^+/TertCI^ mice since the decrease in p21 levels was more pronounced than in the p21^+/-^ mice. It was previously reported that a catalytically inactive TERT was able to cause proliferation of hair follicle (Sarin *et al*, 2005), skin basal keratinocytes (Choi *et al*, 2008) and kidney podocytes (Shkreli *et al*, 2011). In particular, transient overexpression of catalytically inactive TERT (i-TERT^ci^ mouse model) triggers the clonal expansion of podocyte progenitor cells, however these cells are unable to differentiate inducing lethal nephropathy (Montandon *et al*, 2022). Overall, we speculate that the decreased expression of p21 favours the expansion of progenitor cells in both p21^+/Tert^ and p21^+/TertCI^ mice, however only the mice that express catalytically active telomerase maintain this effect during aging by preventing cellular senescence.

Previous study showed that after pneumotectomy, stimulated pulmonary caplillary EC marked with VE-cadherin, VEGFR2, FGFR1, and CD34+ produce angiocrine factors that produce proliferation of EC progenitors supporting alveologenesis (Ding *et al*, 2011) A subclass of endothelial cells interacting with AT1 cells and expressing high levels of Car4, Kdr, and CD34 was further described as essential for regeneration of pulmonary alveolar endothelium after acute lung injury (Niethamer *et al*, 2020). Of note, these cells are primed to respond to VEGFA due to their high level of VEGF receptor^39^. In our study, the single-cell experiments provide a framework to identify the nature of the CD34+ endothelial cells and their properties. We found that gCap cells and aerocytes both expressed CD34 but aerocytes expressed CD34 at highest levels. We report in addition that CD34+ cells in the p21^+/Tert^ mice are marked by higher incorporation of BrdU and higher PCNA labelling reflecting their ability to proliferate. Because aerocytes have been described to rarely proliferate even after injury (Gillich *et al*, 2020), the question remains about the identity of the proliferating CD34+ cells. Because aerocytes may be generated from gCap during repair (Gillich *et al*, 2020), one possibility is that gCaps that express CD34 at low levels generate aCaps marked by high CD34 levels in p21^+/Tert^ mice. This hypothesis is consistent with the fact that we found that the ratio aCap/gCap was higher in p21^+/Tert^ mice compared to the control mice. Alternatively, since both cell types develop from bipotent progenitors (Gillich *et al*, 2020), another possibility is that p21-promoter dependent expression of Tert promotes the differentiation of the progenitors into aerocytes.

Because, we were unable to reveal the differences in the levels of Cd34+ mRNA among the four mouse models by scRNA-seq, in contrast to what we observed in the immunofluorescence experiment revealing the CD34 protein, it may that TERT could regulate CD34 levels at a post-transcriptional level.

In conclusion, our results support a model in which aging mice accumulate senescent endothelial cells with critically short/damaged telomeres. In the p21^+/Tert^ mouse, p21-promoter dependent expression of TERT will favour stem cell mobilization. At the same time, TERT expression will prevent these cells from becoming senescent. This process would be particularly important in maintaining the capillary density. As a consequence, this will reduce age-related emphysema and perivascular fibrosis.

Overall, our results open new avenues for the development of treatments based on the conditional expression of TERT targeting lung endothelial cells to cure lung diseases. Nevertheless, the oncogenic potential of telomerase expression under the control of the p21 promoter will have to be evaluated in a very rigorous way, even if we did not observe in p21+/Tert mice any lung cancer. In the recent years, trials have been made to explore the potential of CD34+ cell therapy for the treatment of different cardio vascular disease (Prasad et al, 2020). We can anticipate that the long-term maintenance of CD34+ endothelial cells could have therapeutic benefits beyond pulmonary pathologies.

## MATERIALS AND METHODS

### mCherry-2A-Tert and mCherry-2A-Tert^CI^ mice generation

Mouse lines generated in this study are deposited at Ciphe, Marseille, France. Generation of the p21^+/Tert^ mouse model. A 6,4 kb genomic fragment encompassing coding exons 2 and 3 of the *Cdkn1a* gene was isolated from a BAC clone of B6 origin (clone n° RP23-412O16; http://www.lifesciences.sourcebioscience.com) and subcloned into the *Not* I site of pBluescript II. The genomic sequence containing the homologous arms was checked by DNA sequencing. A synthetic gene encoding the mCherry, 2A peptide, and the 5’ coding region of mTert (beyond the *Sac* II site) was constructed by gene synthesis (By DNA2.0 Inc) to produce the mCherry-2A-mTert^*Sac*II^ cassette. Using ET recombination, the synthetic mCherry-2A-mTert^*Sac*II^ cassette was introduced in the 5’ coding region of the *Cdkn1a* gene replacing the start codon. The full-length mTert was next introduced in the mCherry-2A-mTert^*Sac*II^ truncated cassette by inserting into the targeting vector a synthetic mTert (*Sac* II-*EcoR* I) fragment restoring the full-length mTert to give rise to the complete mCherry-2A-mTert cassette. A self-deleter NeoR cassette was further introduced in the *EcoR* I site of the targeting vector immediately after the mTert sequence. A cassette coding for the diphtheria toxin fragment A expression cassette was finally introduced in the *Not* I site of the targeting vector. All the elements of the final targeting are shown in the *supplementary dataset 1*.

JM8.F6 C57BL/6N ES cells were electroporated with the targeting vector linearized with Fse1 (Pettitt *et* al, 2009). The scheme of the introduction of the mCherry-2A-mTert-Neo^R^ cassette in place of the ATG codon of Cdkn1a is shown in the source data (Source data-Fig.1A, 1B). After selection with G418, ES cell clones were screened for proper homologous recombination. The correct integration of the cassette was verified by Southern blot with a Cdkn1a probe (Source data Fig.1C) and by long range PCR (Source-data Fig. 1D). The primers used to check the correct insertion of the cassette were:

1310_ScES_01 Fwd: 5’ CTGAATGAACTGCAGGACGA ;

1310_ScES_01_Rev : 5’ CTTGCCTATATTGCTCAAGG.

To ensure that adventitious non-homologous recombination events had not occurred in the selected ES clones, a Southern blot was performed with a mCherry probe (Source data Fig. 1E).

Properly recombined ES cells were injected into Balb/c blastocysts. Germline transmission led to the self-excision of the loxP-Cre-NeoR-loxP cassette in male germinal cells. p21-mCherry-2A-mTert mice (p21^+/Tert^) were identified by PCR of tail DNA. The primers used for genotyping were as follows: sense-WT 5’-GCTGAACTCAACACCCACCT-3’, sense-p21-Tert 5’-GGACCTCTGAGGACAGCCCAAA-3’ and antisense 5’-GCAGCAGGGCAGAGGAAGTA-3’. The resulting PCR products were 435 (WT) and 520 (p21^+/*Tert*^) base pairs long. The proteins produced by the mRNA mcherry-2A-mTert polycistronic mRNA are shown in Source data-Fig 1F. In the final construct, Cdkn1a protein is no further produced from the mutated allele because of the substitution of the C*dkn1a* [ATG] codon by the cassette and the presence of a STOP codon at the end of Tert.

As control mice, we developed p21^+/+^ littermates and p21^+/-^ mice that were obtained by crossing the p21 homozygous knockout strain B6.129S6(Cg)-*Cdkn1a*^*tm1led*^ (from JAX laboratories) with isogenic p21^+/+^ mice. In the p21^+/-^ mouse, a neo cassette replaces exon 2 of *Cdkn1a*. Homozygotes are viable, fertile, and of normal size.

p21^+/*TertCI*^ mice were constructed with exactly the same targeting vector used to construct p21^+/*Tert*^ except that the codon GAT in *Tert* coding for D702 has been changed into GCA coding for A. Primers to genotype p21^+/*Tert*^ and p21^+/*TertCI*^ mice are indicated in Supplemental Information.

### Ethical statement

Mice were used according to institutional guidelines, which complied with national and international regulations. All animals received care according to institutional guidelines, and all experiments were approved by the Institutional Ethics committee number 16, Paris, France (licence number 16-093). Mice were bred and maintained in specific-pathogen-free conditions with sterilized food and water provided ad libitum and were maintained on a 12 h light and 12 h dark cycle in accordance with institutional guidelines.

### Verification of the TERT activity

To check that TERT expressed from the mCherry-2A-mTert can produce active telomerase, the mCherry-2A-mTert was amplified by PCR (S 5’-atatat<u>gaattc</u>atggtgagcaagggcgaggaggata; AS 5’-atatatgcggccgcttagtccaaaatggtctgaaagtct) from the targeting vector and cloned into the *EcoR* I/*Not* I sites of pCAG-GFP. In the resulting vector, the mCherry-2A-mTert cassette is expressed under the control of the strong constitutive CAG promoter. The resulting vector was transfected in the mTert-/-mouse ES cells (provided by Lea Harrington, Montreal). Prior to transfection, the Neo^R^ cassette in the Tert-/-ES cells was disrupted by CRISPR/CAS9 since the Tert-/-ES cells and the electroporated plasmid shared the same antibiotic resistance. The presence of telomerase activity was analyzed in the cell extracts using the Telomere Repeat Amplification Protocol (TRAP) (see below). We could detect robust telomerase activity in mouse ES mTert-/-transfected by the construct pCAG-mCherry-2A-mTert (Fig. 1B).

### Fluorescence imaging of mice and organs

To demonstrate *in vivo* the expression of the KI *Cdkn1a* allele in response to p21 promoter activation, littermate p21^+/+^ and p21^+/Tert^ mice were exposed either to a whole body ionizing radiation or to doxorubicin which both activate the p53-p21 axis of the DNA damage response (DDR). For irradiation experiments, p21^+/+^ and p21^+/Tert^ mice were imaged (time zero) and then exposed to 1.5-Gy of total body irradiation (RX, RS2000 Irradiator, Radsources). Non-invasive whole-body fluorescence (FLI) was determined at the indicated times post-treatment. Imaging was performed using a Photon-IMAGER (Biospace Lab), and mice were anesthetized with 3 % vetflurane through an O_2_ flow modulator for 5 min and then the image was acquired (4-s exposure, excitation=545nm, background=495nm, emission filter cut off=615nm). Corresponding color-scale photographs and color fluorescent images were superimposed and analyzed with M3vision (Biospace Lab) image analysis software.

For Doxorubicin treatments, the p21^+/+^ and p21^+/Tert^ mice were injected i.p. with doxorubicin (DXR) (20 mg/kg). Doxorubicin has been shown to activate p21 promoter to high levels in the liver and kidneys (Tinkum *et* al, 2011). 24h after DXR injection, mice were then sacrificed and organs were rapidly removed and imaged within 15 min of sacrifice (4-s exposure). After that, proteins and RNA were extracted from kidneys and liver for immunoblot (p21) and RT-qPCR (Tert) analyses, respectively.

### Primary pulmonary artery smooth muscle cells and culture

Pulmonary artery smooth muscle (PA-SMC) were extracted and cultured as already described (Kasahara *et* al, 2011). Cells were cultured at 37°C, 5% O2, in DMEM (High glucose, -Pyruvate, Gibco) supplemented with 10%(v/v) decomplemented FBS and 1%(v/v) Penicillin-Streptomycin. At each passage, cells were counted and 50,000 cells were put back in culture in a 25 cm2 flask and cultured in DMEM supplemented with FBS. Representative images of cells in culture were obtained using EVOS M5000 Imaging System (ThermoFisher) on p21^+/+^ and p21^+/Tert^ cells 24h after plating on a 24 well plate.

### Animal studies, lung tissue analysis

Groups of mice from the four genotypes were prepared in order to sacrifice 4-month-old mice and 18-month-old mice. To reach the final mice number per group and per genotype, this experiment was reproduced three individual times, on three independent cohorts. For p21^+/Tert-CI^ mice, an independent cohort of p21^+/Tert^ and p21^+/Tert-CI^ mice was prepared.

A subgroup of mice received an intraperitoneal injection of 5-bromo-2’-deoxyuridine (ab142567, abcam) diluted in PBS at 25mg/ml. Mice received 100mg/kg BrdU 24h prior to sacrifice. Mice were anesthetized with an intraperitoneal injection of ketamine (60 mg/kg) and xylazine (10 mg/kg). After mice sacrifice three lobes of the right lung were quickly removed and immediately snap-frozen in liquid nitrogen then stored at -80°C for biological measurements. Genomic DNA from the right lung of each animal was used for TeSLA assay. The last lobe of the right lung was fixed with 2% formaldehyde (Sigma) and 0.2% glutaraldehyde (Sigma) for 45 minutes. Then, lungs were washed with PBS and stained overnight in a titrated pH 6 solution containing 40 mM citric acid, 150 mM NaCl, 2 mM MgCl_2_, 5 mM potassium ferrocyanide, and 1 mg/ml X-Gal at 37°C. Stained lobes were then imbedded in paraffin and 5μm thick sections were cut. Lung sections were deparaffinized using xylene and a graded series of ethanol and then nuclei were stained with neutral red. 10 fields per section were acquired at an overall magnification of 500. The number of beta-galactosidase stained cells and expressed as a % of total number of cells per field.

The left lungs were fixed by intratracheal infusion of 4% paraformaldehyde aqueous solution (Sigma) at a trans-pleural pressure of 30 cm H_2_O and processed for paraffin embedding. For morphometry studies, 5 μm–thick sagittal sections along the greatest axis of the left lung were cut in a systematic manner to allow immunostaining. Lung emphysema was measured using mean linear intercept methods on hematoxylin-eosin-safran (HES) coloration. Briefly 20 fields/animal light microscope fields at an overall magnification of 500, were overlapped with a 42-point, 21-line eyepiece according to a systematic sampling method from a random starting point. To correct area values for shrinkage associated with fixation and paraffin processing, we used a factor of 1.22, calculated during a previous study (Houssaini *et* al, 2018). In addition, MLI measurement was performed by an independent operator by using semiautomated measurement of MLI (Crowley *et al*, 2019). Both methods (manual and semi-automated) performed by two independent observers provided the same results.

Lung fibrosis was quantified on lung sections stained with Sirius Red (Picro Sirius Red Stain Kit, ab150681, Abcam, Cambridge, UK) using the modified Ashcroft scale (Hübner *et al*, 2008). Briefly, the lungs were scanned microscopically with a 20-fold objective, which allowed the evaluation of fine structures while also providing a sufficiently broad view. The entire section was examined by inspecting each field in a raster pattern. Areas with dominating tracheal or bronchial tissue were omitted. The grades were summarized and divided by the number of fields to obtain a fibrotic index for the lung.

### Immunofluorescence

Paraffin-embedded sections of lung were deparaffinized using xylene and a graded series of ethanol dilutions then processed for epitope retrieval using citrate buffer (0.01 M, pH 6; 90°C, 20 min). For CD31 epitope retrieval, we used Antigen retrieval buffer (100X Tris-EDTA buffer, pH9.0, ab93684, Abcam) according to the manufacturer’s instructions. For nuclear immunolabeling, tissues were permeabilized with 0.1% Triton X-100 in PBS for 10 minutes. Saturation was achieved using Dako antibody diluents with 10% goat serum. For immunolabelling with primary antibody produced in mouse we used the M.O.M. (mouse on mouse) immunodetection kit, basic (ref. BMK-2202, Vector Lab) according to manufacturer instructions. For double staining, first and second primary antibodies were diluted in Dako antibody diluents with 3% goat serum then incubated for 1 hour at 37°C in a humidified chamber. After PBS washes, the sections were covered with secondary antibody (Dako antibody diluents with 3% of goat serum mixed with mouse or rabbit Alexa Fluor® 555 or Alexa Fluor® 660 (Termofisher)) for 40 minutes at 37°C in a humidified chamber. The sections were secured with fluorescent mounting medium containing DAPI and protected with coverslips. Fluorescence was recorded using an Axio Imager M2 imaging microscope (Zeiss, Oberkochen, Germany) and analysed on digital photographs using Image J software (imagej.nih.gov/ij/). Following primary antibodies were used: rabbit anti-CD31 (ab182981, Abcam, 1:500); rabbit anti-MUC1 (ab109185, Abcam, 1:200); rabbit anti-CD34 (ab81289, Abcam, 1:200); mouse anti-CDKN2A/p16INK4a (ab54210, 1:200); mouse anti-PCNA (proliferating cell nuclear antigen) (ab29, Abcam, 1:200); mouse anti-BrdU (ab8152, Abcam, 1:20). BrdU immunohistostaining was performed by using BrdU immunohistochemistry kit following provider instruction (ab125306, Abcam).

### RNA extraction, RT-qPCR, and Western blots

Total RNA was extracted using the method (Chomczynski & Sacchi, 1987) with reagents included in the RNeasy kit (Qiagen). To exclude contamination with genomic DNA, the RNA was treated with DNase I directly on mini columns. Reverse transcription was performed using 0.5 to 2.0 μg of RNA, 50 ng/μL random hexamers, and 200 U of Superscript IV (Invitrogen) in the 20 μL reaction volume for 15 min at 55°C, followed by inactivation for 10 min at 80°C. The resulting cDNA was diluted 5-20 fold and analyzed by real-time qPCR using the SYBR Green master mix (Takara bio) and 400 nanomolar each of the following primers: mTert-2253S 5’-AGCCAAGGCCAAGTCCACAA and mTert-2399A 5’-AGAGATGCTCTGCTCGATGACA to target all *Tert* transcripts/isoforms; 2A-F2 5’-AGCAGGAGATGTTGAAGAAAACCC and mTert-5’R25’-GGCCACACCTCCCGGTATC to target *mCherry*-*2A*-*Tert* transcript from the KI *Cdkn1a* allele; mActb-90S 5’-ACACCCGCCACCAGTTCG and mActb-283A 5’-GGCCTCGTCACCCACATAGG to target β actin transcript. At least one primer within each pair was designed to anneal onto the exon-exon junction to avoid priming on genomic DNA, and the specificity of amplification was verified on agarose gel.

Immunoblots were carried out using standard procedures. The membranes were incubated with primary antibodies followed by incubation with the secondary HRP conjugates, and the signal was detected using an enhanced chemiluminescence detection system (GE Healthcare). The unsaturated images were acquired using ChemiDoc MP imaging system (Bio-Rad), and the signal densities were quantified using Image Studio Lite ver 5.2 (LI-COR). Antibodies used were rabbit monoclonal anti-p21 [EPR3993] (abcam) and mouse anti-b-actin (SigmaAldrich A1978) antibody.

### Telomere Repeat Amplification Protocol (TRAP)

To measure telomerase activity in cellular/tissue extracts TRAP was performed according to the original protocol (Kim & Wu, 1997; Kim *et al*, 1994). Briefly, the cells or tissues were extracted on ice using standard CHAPS buffer (60 μL per 10^6^ cells) supplemented with a cocktail of protease inhibitors (Roche) and 20 units of RNase inhibitor (Applied Biosystems) followed by centrifugation at 18000 x g for 30 min at 4°C. The protein concentration in the supernatant was quantified using Pierce™ 660nm Protein Assay and the aliquots equivalent to 800, 400, and 200 ng of protein were used to extend 50 ng of the telomerase substrate (TS oligo) in the 25 μL reaction volume. For gel-based detection of the TRAP product we used FAM-labeled TS and the standard ACX and NT primers (Kim *et al*, 1994) to amplify the telomerase extension products and the internal TSNT control, respectively. For the real-time qPCR-based TRAP we used unlabeled TS and the ACX primer alone. The qTRAP was performed using the SYBR Green PCR mix, and the amount of CHAPS extracts was optimized by serial dilutions.

### Telomere Shortest Length Assay (TeSLA)

TeSLA was performed according to the protocol described by Lai *et al*, 2017. Briefly, 50 ng of undigested genomic DNA was ligated with an equimolar mixture (50 pM each) of the six TeSLA-T oligonucleotides containing seven nucleotides of telomeric C-rich repeats at the 3′ end and 22 nucleotides of the unique sequence at the 5’ end. After overnight ligation at 35°C, genomic DNA was digested with *Cvi*AII, *Bfa*I, *Nde*I, and *Mse*I, the restriction enzymes creating either AT or TA overhangs. Digested DNA was then treated with Shrimp Alkaline Phosphatase to remove 5′ phosphate from each DNA fragment to avoid their ligation to each other during the subsequent adapter ligation. Upon heat-inactivation of phosphatase, partially double-stranded AT and TA adapters were added (final concentration 1 μM each) and ligated to the dephosphorylated fragments of genomic DNA at 16°C overnight. Following ligation of the adapters, genomic DNA was diluted to 20 pg/μL, and 2-4 μL was used in a 25 μL PCR reaction to amplify terminal fragments using primers complementary to the unique sequences at the 5’ ends of the TeSLA-T oligonucleotides and the AT/TA adapters. FailSafe polymerase mix (Epicenter) with 1× FailSafe buffer H was used to amplify G-rich telomeric sequences. Entire PCR reactions were then loaded onto the 0.85% agarose gel for separation of the amplified fragments. To visualize telomeric fragments, the DNA was transferred from the gel onto the nylon membrane by Southern blotting procedure and hybridized with the ^32^P-labeled (CCCTAA)_3_ probe. The sizes of the telomeric fragments were quantified using TeSLA Quant software (Lai *et al*, 2017).

### Statistical Analysis

Basic statistics, tests for the differences between the means, and one-way analysis of variance (ANOVA) were performed with the GraphPad Prism 9 Software. ANOVA followed by Bonferroni multiple comparison test was used to compare the means of more than two independent groups. Pairwise correlation and multiple regression analyses were performed using Statistica 13.0 software package (StatSoft/Dell).

### Single cell RNAseq

#### Isolation of lung single cells from 18 month-old p21^+/+^ (1x), p21^+/-^(3x), p21^+/Tert^ (5x), and p21^+/Tert^ (2x) mice

Mouse trachea is injected with 1,5ml dispase 50 U/ml (Corning #354235) followed by 0,5 ml agarose 1%. Lung is resected and minced in 3 ml DPBS 1x CaCl^2+^ and MgCl^2+^ (Gibco, 14040-091). Collagenase/dispase 100 mg/ml (Roche #11097113001) is added to the chopped tissue and placed on a rotator at 37 °C for 30 min. Enzymatic activity is inhibited by adding 5 ml of DPBS 1x (Gibco, 14190-094) containing 10% FBS and 1mM EDTA (Sigma #E7889). Cell suspension is filtered through 100 μm nylon cell strainer (Fisher Scientific #11893402), treated by DNase I (Sigma D4527) and filtered again through a 40 μm nylon cell strainer (Fisher Scientific #11873402). Red blood cells are removed by red blood cell lysis (Invitrogen, 00-4333-57) treatment for 90s at room temperature. Finally, lung cells are centrifuged at 150*g*, 4 °C for 6 min, resuspended in DPBS 1x containing 0,02% BSA (Pan Biotech #P06-13911000) and counted in a Malassez.

#### Preparation of single cell sequencing libraries

Single-cell 3’-RNA-Seq samples were prepared using single cell V2 reagent kit and loaded in the Chromium controller according to standard manufacturer protocol (10x Genomics, PN-120237) to capture 6.000 cells. Briefly, dissociated lung cells are encapsulated using microfluidic device. RNAs are captured on beads coated of oligos containing an oligo-dTTT, UMIs and a specific barcode. After reverse transcription, cDNAs are washed, PCR-amplified and washed again before analysis on a Bioanalyzer (Agilent) for quality control. Finally, libraries are prepared following standard Illumina protocol and sequenced on a NovaSeq sequencer (Illumina). Raw sequences are demultiplexed and reads are mapped onto the mm10 reference genome using the v2.3 Cell Ranger pipeline (10X Genomics) to generate a count matrix for each sample.

#### Quality control and Normalization of the single-cell data

The digital matrices were filtered by cell type (on clusters composed of the same cell type), to remove low-quality cells with low UMI counts and cells with relatively high mitochondrial DNA content. Outlier analysis was performed with perCellQCMetrics from the scatter package. An upper cutoff was manually determined for each sample based on a plot of gene count versus UMI count or % of mitochondrial genes, to have at least 1000 UMIs, number of transcripts ranging between 1000 and 30000 and at most 14% mitochondrial transcripts. The quality was consistent across samples, and differences in RNA and gene content could be ascribed to cell-type-specific effects. DGE matrices from all samples, sequenced at different time, were then merged and subsequently normalized using the deconvolution normalization method in the *scran R* package in order to correct for differences in read depth and library size inside and between samples.

#### Clustering, cell type, and cycle annotation

Seurat v4.1.0 was used to perform dimensionality reduction, clustering, and visualization on the unique combined (with all samples) and normalized matrix. After scaling the data, dimensionality reduction was performed using PCA on the highly variable genes. Seurat’s *FindNeighbors* function was run to identify cluster markers with the following parameters: reduction = “pca” and dims = 1:10, followed by the *FindClusters* function with resolution = 1. The Seurat function *RunUMAP* was used to generate 2-dimensional umap projections using the top principal components detected in the dataset. To annotate cells, we mapped each cluster to the Mouse Cell Atlas (Kalucka *et al*, 2020) by using the *scMCA* function (Sun *et al*, 2019). Cluster identities were further verified according to gene markers found with the *FindMarkers* function from Seurat. Doublet cells were identified manually as expressing markers for different cell types and were removed from the matrix. In order to define endothelial (EC) subclusters, EC cell classes were isolated and subjected to a new clustering by the *FindNeighbors* function with new parameters: reduction = “pca” and dims (1:15 for EC). Seurat *FindAllMarkers* function was applied to each EC subclusters and the markers were compared to the top 50 gene markers previously found in lung EC (Gillich *et al*, 2020). We thus annotated EC subclusters depending on the enrichment of marker genes and identified five subtypes of EC (artery, capillary_1, capillary_2, vein and lymphatic). We annotated two functionally divergent lung EC capillary clusters according to Matmati *et al*, 2020. We thus reassign EC capillary_1 as “aCap” and EC capillary_2 as “gCap”. In order to assign cell cycle phase to each cell, we used the Cyclone method to our single cell RNA-Seq dataset.

### Data and materials availability

Original data for single cell RNA-sequencing is available at the Gene Expression Omnibus (GEO), GSE165218.

## ACKNOWLEDGMENTS

We thank Lea Harrington for providing mTert-/-mouse ES cells. We are very grateful to Myriam Grunewald (Hebrew University of Jerusalem) for critically reading the manuscript. We are grateful to the CRCM animal facility for taking care of the mouse strain colonies, to Manon Richaud from the CRCM flow cytometry and cell-sorting platform, and to Lionel Spinelli and Arnaud Guille for useful discussions and technical support for the single-cell analysis. We thank the Institut Curie Bioinformatics platform for data management and quality control of single-cell data.

Work in VG’s Lab is supported by “La Ligue Contre le Cancer”, Equipe Labellisée, La Région SUD (Volet Général), The Canceropole PACA (Projet Emergent), the “Institut National du Cancer” (INCA), PLBIO 2019 and the cross-cutting Inserm program on aging (AgeMed).

SA’s Lab is supported by grants from the INSERM, Ministère de la Recherche, Agence Nationale pour la Recherche (ANR), Institut National du Cancer (INCA), Fondation pour la Recherche sur la Cancer (ARC) and Fondation pour la Recherche Médicale (FRM).

ALV’s lab is supported by grants from ANR (Lustra), INCa (PLBIO2019), EDF (CT9818) and La Ligue contre le cancer-Paris (RS21/75-24). SCA is a recipient of a European CO-FUND PhD fellowship from Institut Curie (European Union’s Horizon 2020 research and innovation programme under the Marie Skłodowska-Curie grant agreement No 666003).

IF’s lab was funded by grants from the Spanish Ministry of Science and Innovation (PID2019-110339RB-I00) and the Comunidad de Madrid (S2017/BMD-3875).

EG’s lab was supported by the Fondation ARC (Program ARC), the ANR grants TELOPOST and the cross-cutting Inserm program on aging (AgeMed).

CIPHE is supported by PHENOMIN (French National Infrastructure for mouse Phenogenomics; ANR10-INBS-07).

The imaging studies carried out on the TrGET platform (IPC/CRCM) received financial support from ITMO Cancer as part of the 3rd Cancer Plan for the acquisition of dedicated equipment.

High-throughput sequencing was performed by the ICGex NGS platform of the Institut Curie supported by the grants ANR-10-EQPX-03 (Equipex) and ANR-10-INBS-09-08 (France Génomique Consortium) from the Agence Nationale de la Recherche (“Investissements d’Avenir” program), by the ITMO-Cancer Aviesan (Plan Cancer III) and by the SiRIC-Curie program (SiRIC Grant INCa-DGOS-465 and INCa-DGOS-Inserm_12554).

## AUTHOR CONTRIBUTION

Conceptualization, M.B., L.L., S.A., A.L.V., I.F. and V.G; Methodology, M.B., L.L., D.C., C.C., C.F., S.C.A., S.B., F.J., L.B., F.F., R.C., E.J., C.S.F., G.G.; Investigation and validation, M.B., L.L., D.C., C.C., C. F., C.S.F., G.G; I.F., E. G., A. L. V., S. A., V.G.; Supervision, M.B., L.L., I.F., A.L.V, S.A., V.G, Project administration, S.A., V.G.; Funding acquisition, I.F., E.G., A.L.V., S.A., V.G, Writing-original draft, VG; Writing Review-Editing, A.L.V., L. L., S.A.

## EXPANDED VIEW FIGURE LEGENDS

**Fig. EV1.**
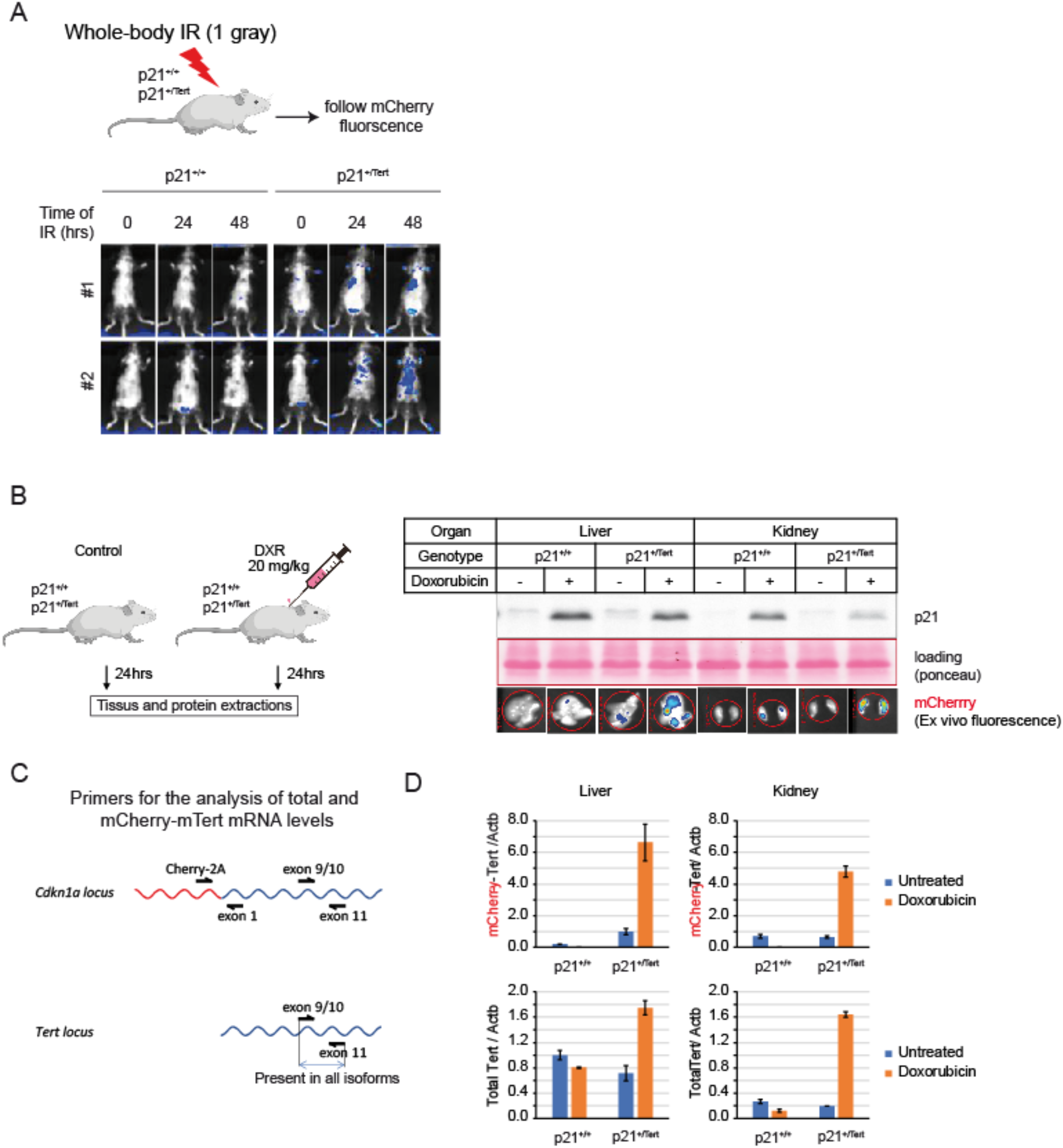
Validation of p21^+/Tert^ model. **(A)** p21^+/+^ and p21^+/Tert^ littermates were subjected to whole-body ionising radiation (1.5 gray). The fluorescence emitted by the mCherry was followed post-irradiation at the indicated times by *in vivo* mCherry imaging (Excitation = 545 nm, Background = 495 nm, Emission = 615 nm). **(B)** Left panel, p21 expression and mCherry fluorescence were analyzed in the liver and kidneys after doxorubicin treatments. Right panel, livers and kidneys of p21^+/+^ and p21^+/Tert^ mice were harvested 24 hours after doxorubicin treatment. The level of p21 protein was evaluated by semi-quantitative immunoblotting and liver and kidney were imaged for mCherry fluorescence. **(C)** Primers to distinguish transgene expression from totat Tert expression are shown **(D)** *Tert* RT-qPCR experiments were performed using RNA extracted from the harvested organs.

**Fig. EV2.**
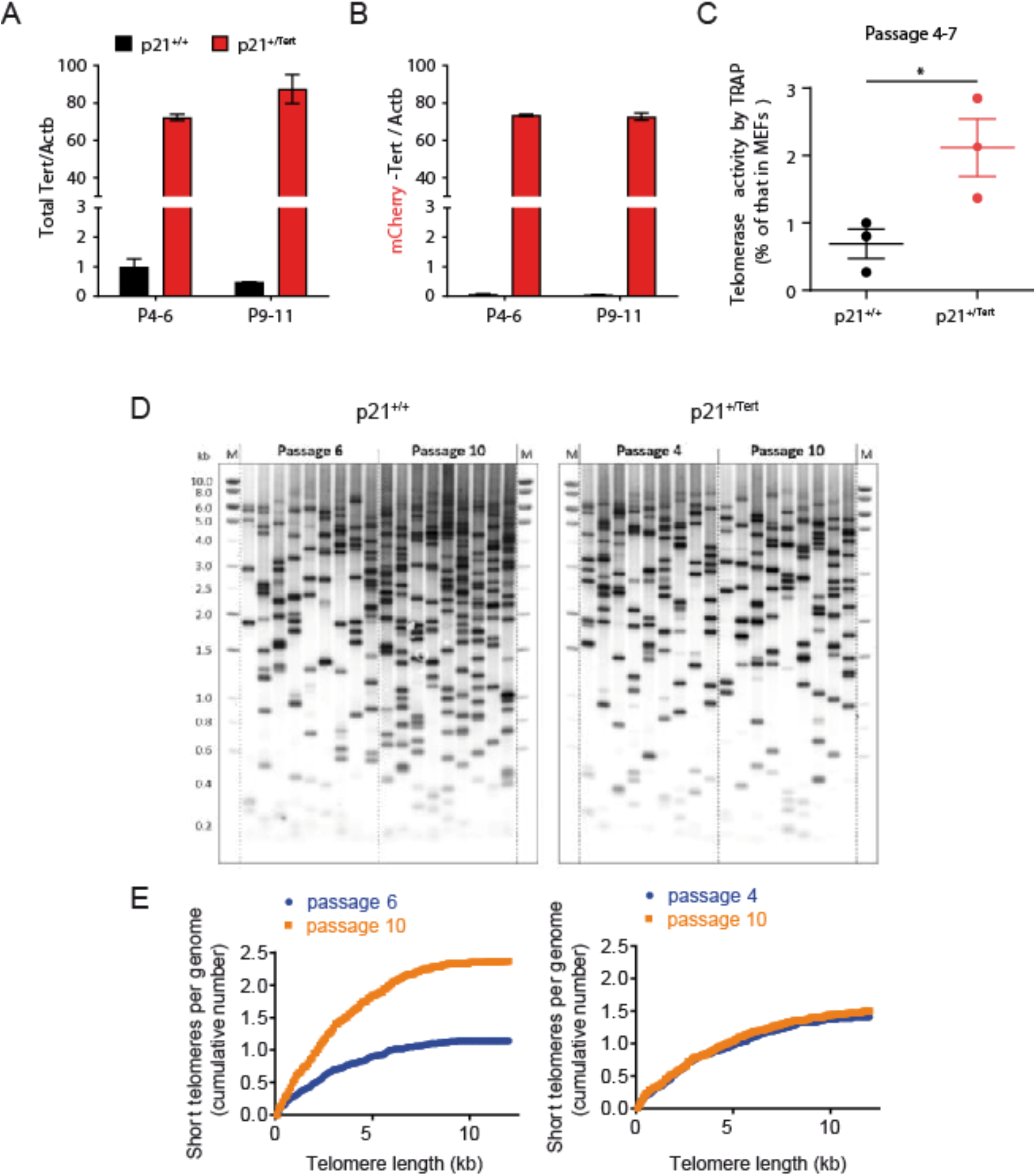
(related to Fig. 1C). p21-promoter dependent TERT expression bypasses senescence in PA-SMCs *ex-vivo* (**A** and **B**) Quantification of the *Tert* mRNA levels in PA-SMCs from the p21^+/+^ and p21^+/Tert^ 4-month-old mice (littermates) at early (p4-6) and late (p9-11) passage. Left panel represent the level of endogenous Tert mRNA, right panel represents the level of KI allele. Nearly all *Tert* mRNA is transcribed from the KI allele. The means of three independent measurements are plotted, and the error bars are SEs. **(B)** Telomerase activity measured by qTRAP at early passages. The data points correspond to vascular PA-SMC cultures established from individual p21^+/+^ and p21^+/Tert^ 4-month-old mice. * p <0.05 from the two-sided *t* test. **(C)** Analysis of the short telomere fraction by Telomere Shortest Length Assay (TeSLA) in the cultured PA-SMCs from p21^+/+^ and p21^+/Tert^ mice. Left panels depict Southern blots probed for the TTAGGG repeats, and the right panels show quantification of the cumulative number of short telomeres across the telomere length thresholds.

**Fig. EV3.**
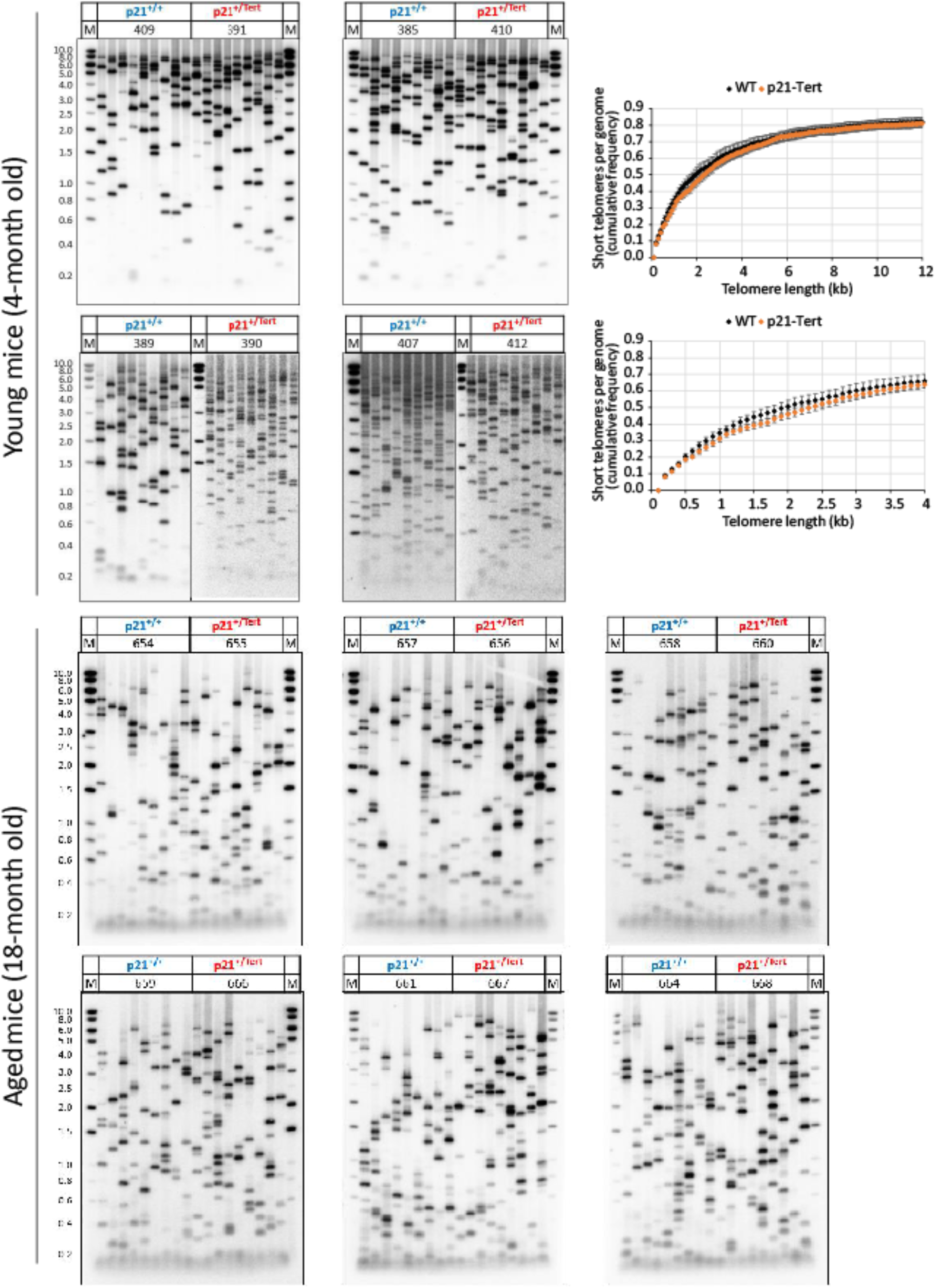
TeSLA Southern blots used for quantification of the Very Short Telomeres in the lungs of the young and old mice. Number of short telomeres per genome in lungs from the p21^+/+^ and p21^+/Tert^ littermates (4 and 18-month-old mice). Genomic DNA was extracted from whole lungs and the length of short telomeres; TeSLA was initiated with 50 ng of genomic DNA extracted from mouse lungs. In the final PCR step, 500 pg of ligation product was used per reaction, and 9 independent reactions were performed for each sample (each lane correspond to an independent PCR reaction) to achieve > 100 amplified telomeres for quantification. Note that young mice, regardless of their genotype, have less telomeres shorter than 1 kb compared to the old mice.

**Fig. EV4.**
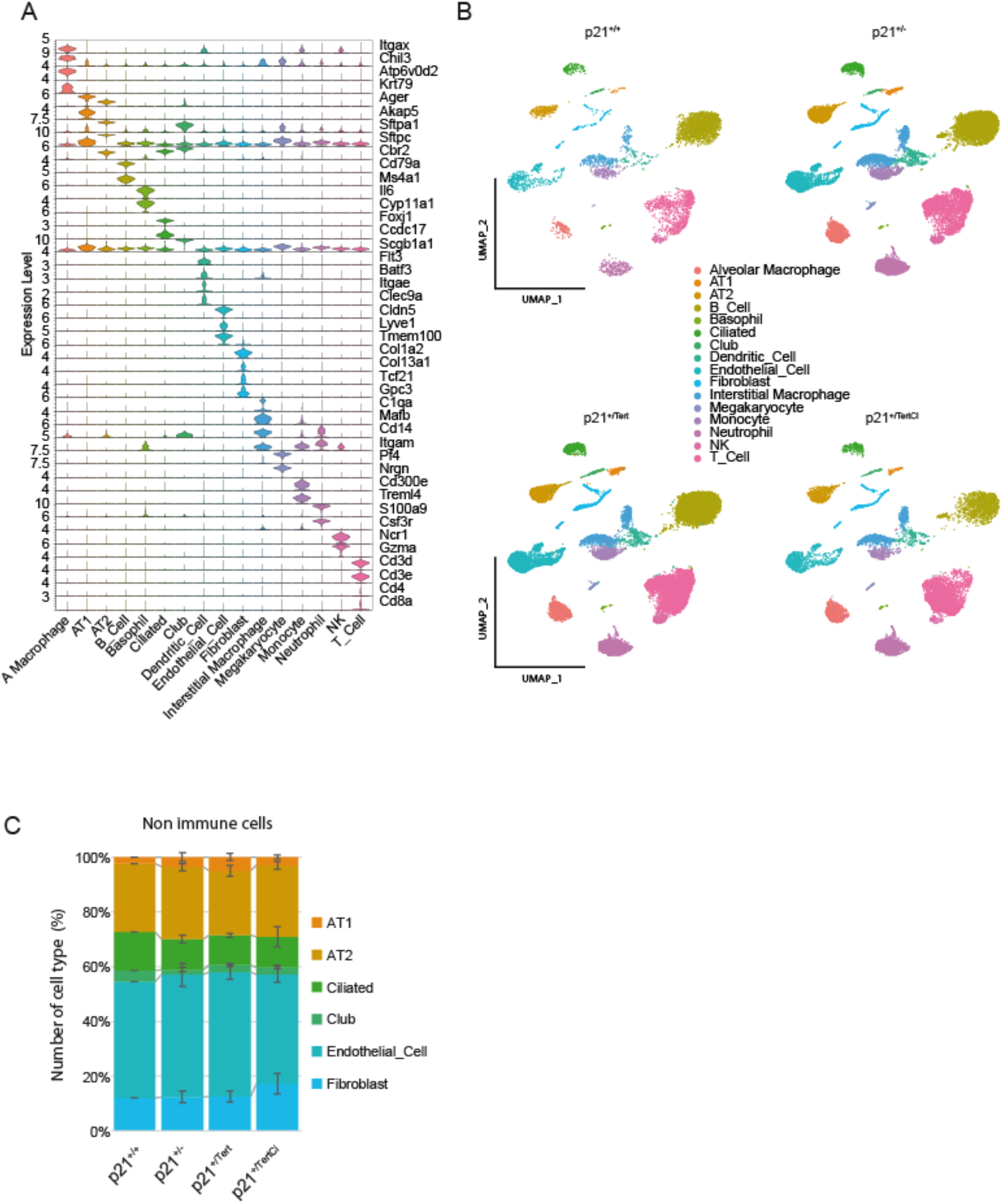
Lung cell types in the four mouse models. **(A)** Representative markers use to annotate lung cell types in the four mouse models. **(B)** UMAP clustering of lung cells. Lung cell populations were identified in lung samples from WT (p21^+/+^), p21^+/-^, p21^+/Tert^ and p21^+/TertCI^ 18 month-old mice. **(C)** Distribution on lung cell types in lungs of the 4 genotypes.

**Fig. EV5.**
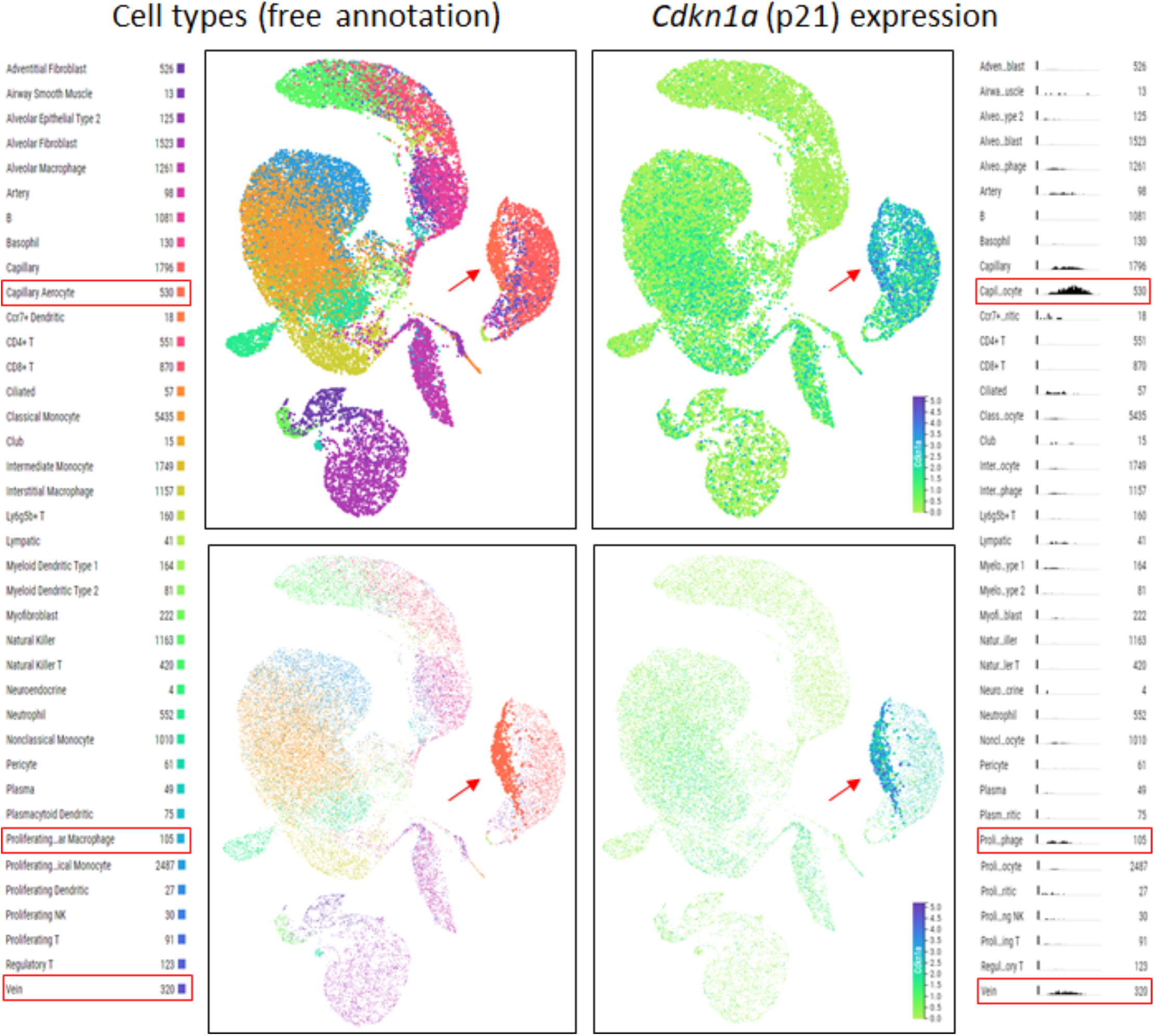
UMAP plots generated using lung droplet scRNA-seq data in Tabula Muris Senis. The data for mice of all ages (1-30 months) were included. In the top panels all cell types are shown, while in the bottom panels only the capillary aerocyte population is highlighted. Red arrow points to the capillary aero-cytes, and red boxes in the annotation mark cell types with elevated p21 expression level. Histograms (on the right) show scaled number of cells expressing p21 (y axis) versus expression level (x axis).

**Fig. EV6.**
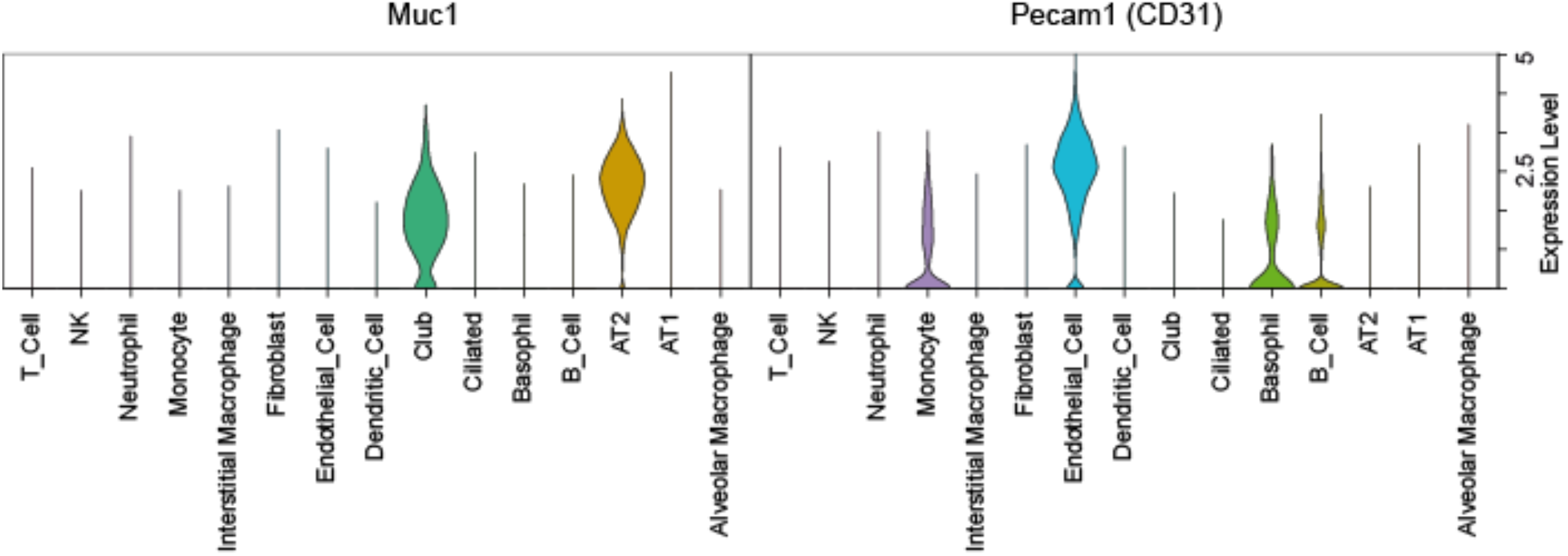
Pecam1 (Cd31) and Muc1 expression in lung cell types.

**Fig. EV7.**
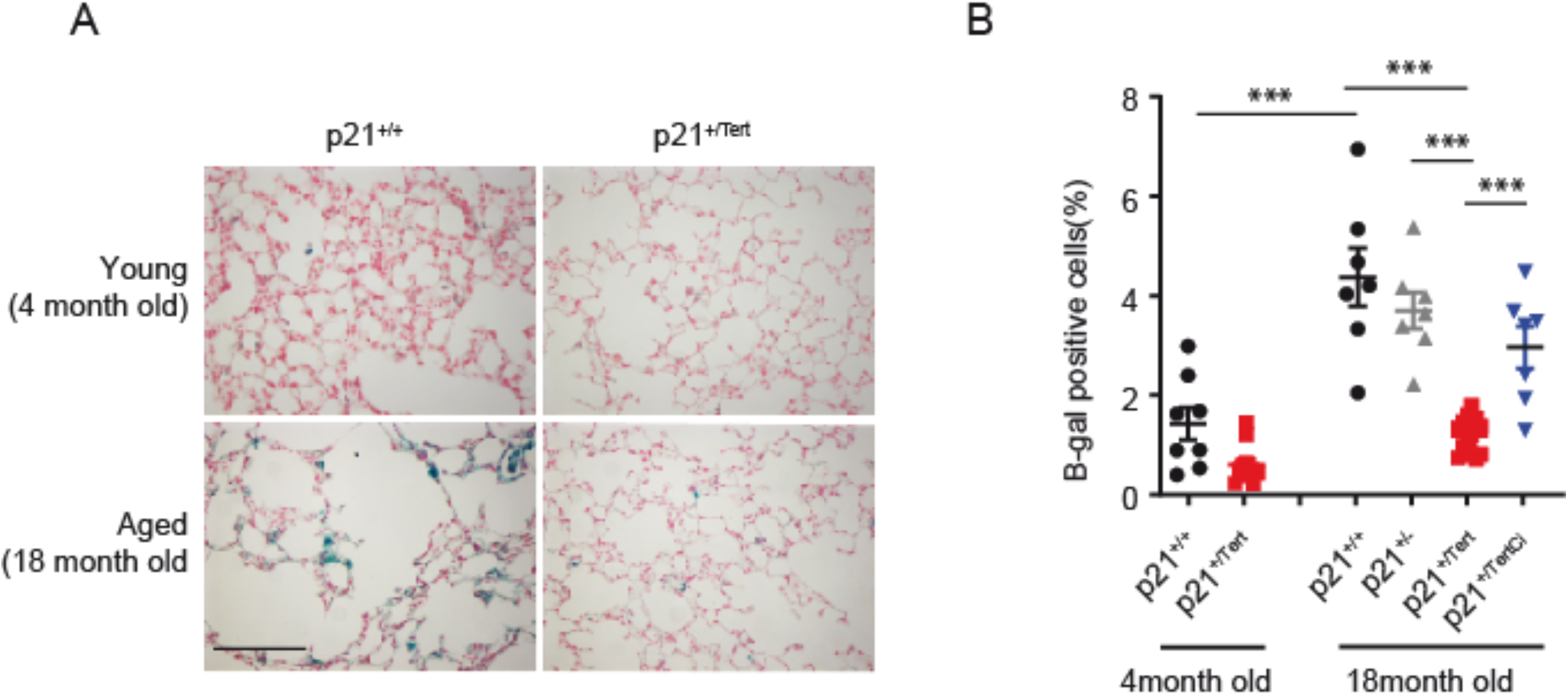
p21^+/Tert^ mice are protected against age-related associated senescent cells accumulation in the lung. **(A)** Representative micrographs of SA-b-Gal staining (blue) in the lungs of young (4-mo-old) and aged (18-mo-old) p21^+/+^ and p21^+/Tert^, Red: fast red nuclear staining. Bar: 50μ. **(B)** Scatter-plot graph showing the percentage of SA-b-Gal positive cells in each group. ***P<0.01 (one way ANOWA followed by Bonferroni post hoc test).

**Fig. EV8.**
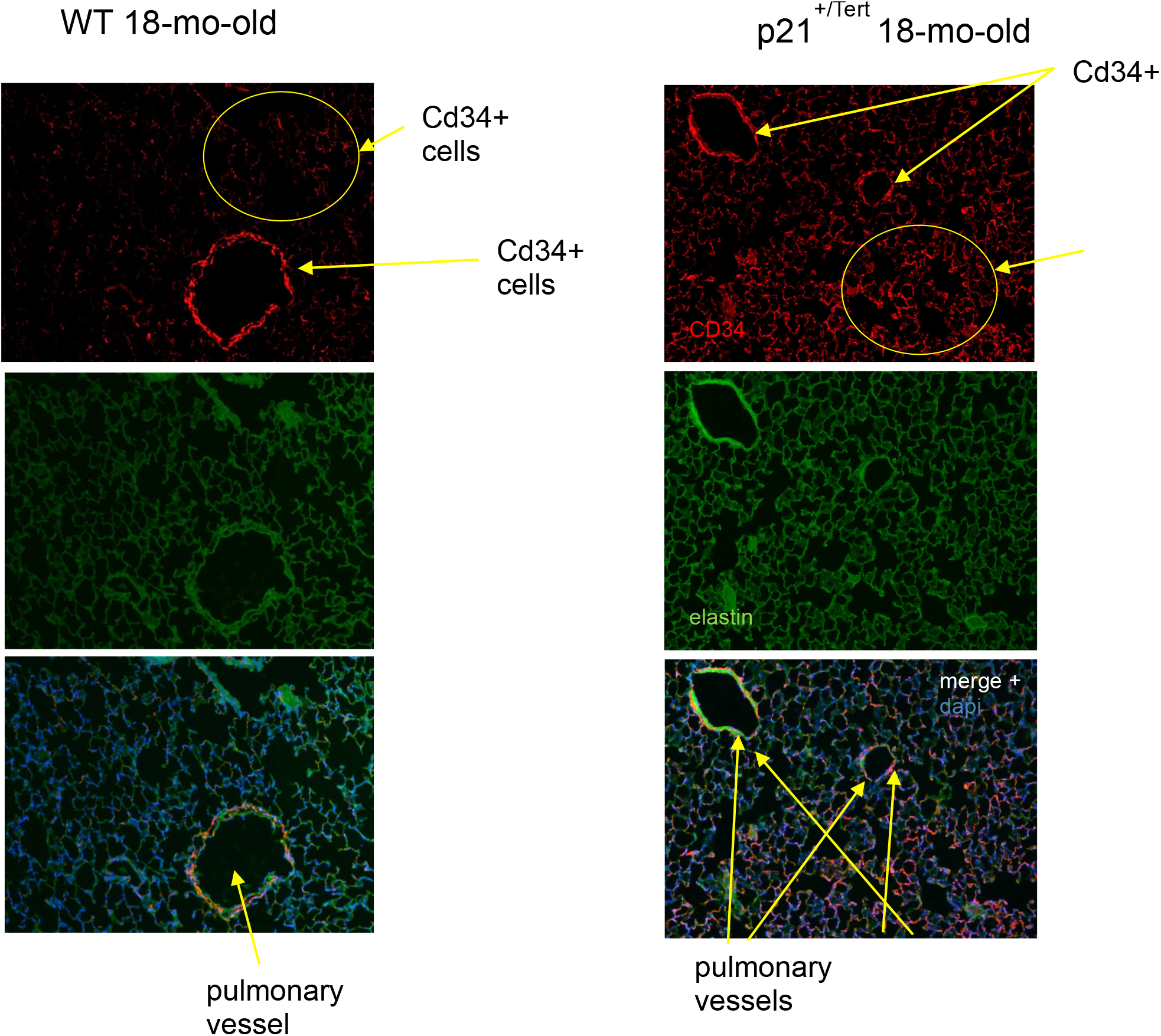
Representative micrographs showing immunofluorescence of CD34 (related to Fig. 6). Cd34 (Red), DAPI nuclear staining (blue). Green elastin (Magnification 10X).

**Fig. EV9.**
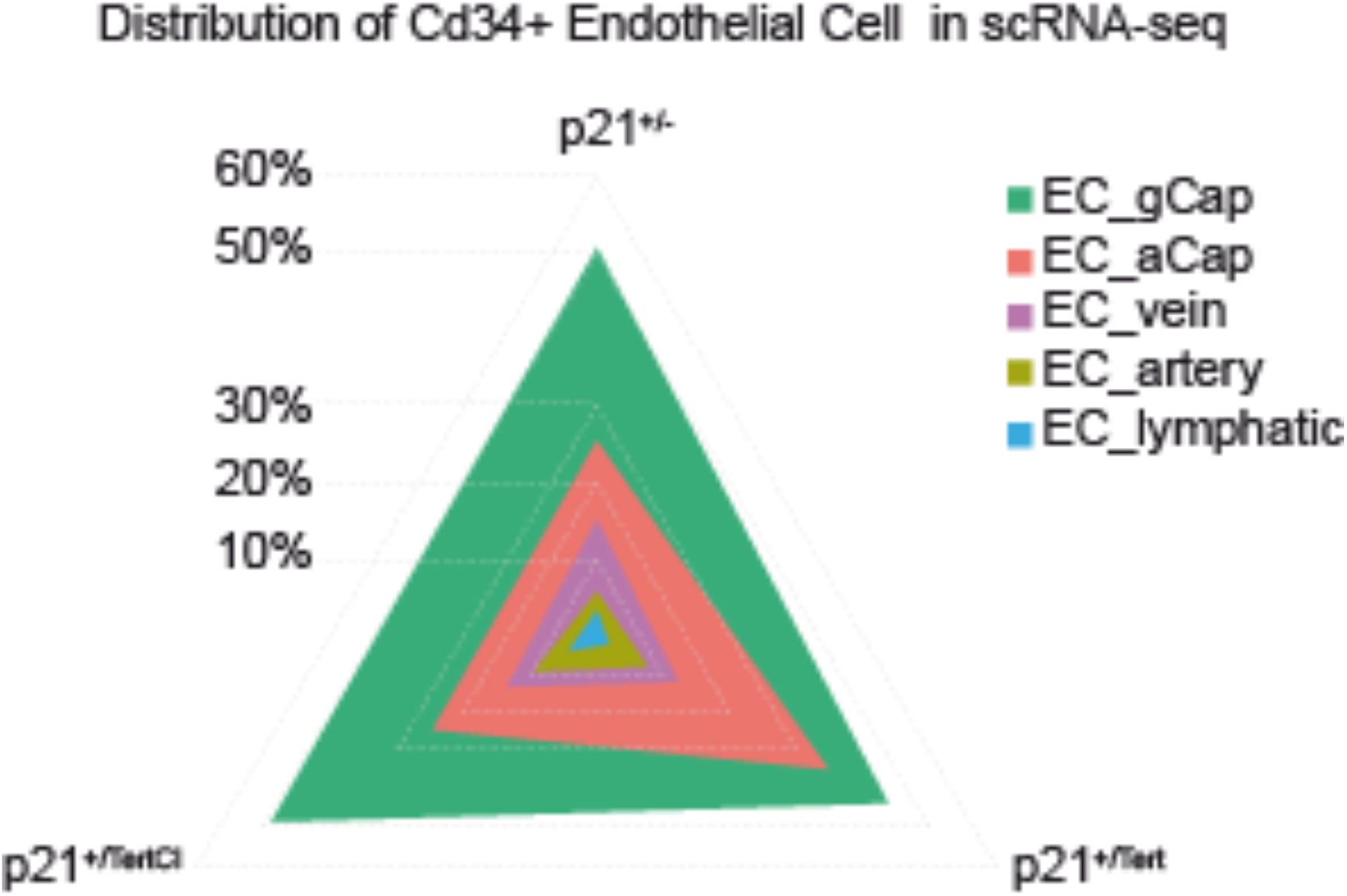
Distribution of Cd34+ (log(counts) > 0.1) in EC subtypes in p21^+/-^, p21^+/Tert^ and p21^+/TertCI^ mice. The number of cells per subtype is expressed as a percentage of the total number of Cd34+ ECs.

**Table EV1**. Number and % of cell types analyzed in the single cell experiments. Refers to Fig. EV4

**Figure source data**. Construction of the p21^+/Tert^ mouse model.

## REFERENCES

Abbas T, Dutta A (2009) p21 in cancer: intricate networks and multiple activities. Nat Rev Cancer. 6:400–14

Alder J K et al. (2015) Telomere dysfunction causes alveolar stem cell failure. Proc Natl Acad Sci 112: 5099–5104

Alder J K et al. (2011) Telomere length is a determinant of emphysema susceptibility. Am J Respir Crit Care Med 184: 904–912

Amsellem V et al (2011) Telomere dysfunction causes sustained inflammation in chronic obstructive pulmonary disease. Am J Respir Crit Care Med 184: 1358–1366

Armanios M & Blackburn E H (2012) The telomere syndromes. Nat Rev Genet 13: 693–704

Armanios M Y et al. (2007) Telomerase mutations in families with idiopathic pulmonary fibrosis. N Engl J Med 356: 1317–1326

Barnes P J, Baker J, Donnelly L E (2019) Cellular Senescence as a Mechanism and Target in Chronic Lung Diseases. Am J Respir Crit Care Med 200(5):556–564

Bernardes de Jesus B et al (2012) Telomerase gene therapy in adult and old mice delays aging and increases longevity without increasing cancer. EMBO Mol Med 4: 691–704

Birch J et al (2015) DNA damage response at telomeres contributes to lung aging and chronic obstructive pulmonary disease. Am J Physiol Lung Cell Mol Physiol 309: L1124–1137

Born E et al (2022) Eliminating senescent cells can promote pulmonary hypertension development and progression. Circulation (in press)

Campisi J (2013) Aging, cellular senescence, and cancer. Annu Rev Physiol 75: 685–705

Chen R et al (2015) Telomerase Deficiency Causes Alveolar Stem Cell Senescence-associated Low-grade Inflammation in Lungs. J Biol Chem 290: 30813–30829

Cheng T et al (2000) Hematopoietic stem cell quiescence maintained by p21cip1/waf1. Science 287 (5459):1804–1808

Childs B G, Durik M, Baker D J & van Deursen J M (2015) Cellular senescence in aging and agerelated disease: from mechanisms to therapy. Nat Med 21: 1424–1435

Choi J et al (2008) TERT promotes epithelial proliferation through transcriptional control of a Myc- and Wnt-related developmental program. PLoS Genet 4(1):e10

Chomczynski P & Sacchi N (1987) Single-step method of RNA isolation by acid guanidinium thio-cyanate-phenol-chloroform extraction. Anal Biochem 162: 156–159

Cordasco E, Beerel F, Vance J, Wende R & Toffolo R R (1968) Newer Aspects of the Pulmonary Vasculature in Chronic Lung Disease. Angiology 19: (7):399–407

Crowley G, Kwon S, Caraher E J, Haider S H, Lam R, Batra P, Melles D, Liu M, Nolan A (2019) Quantitative lung morphology: semi-automated measurement of mean linear intercept. B M C Pulm Med 9: 19(1): 206

Ding B S et al. (2011) Endothelial-derived angiocrine signals induce and sustain regenerative lung alveolarization. Cell 147(3):539–53

Dunnill M S (1962) Quantitative Methods in the Study of Pulmonary Pathology. Thorax 17: 320–328

Fouquerel E et al (2019) Targeted and Persistent 8-Oxoguanine Base Damage at Telomeres Promotes Telomere Loss and Crisis. Mol Cell 75: 117–130 e6

Gillich A et al (2020) Capillary cell-type specialization in the alveolus. Nature 586: 785–789

Grunewald M et al (2021) Counteracting age-related VEGF signaling insufficiency promotes healthy aging and extends life span. Science 30: 373 (6554)

Han X et al (2018) Mapping the Mouse Cell Atlas by Microwell-Seq. Cell 172: 1091–1107 e17

Harper J W et al. (1995) Inhibition of cyclin-dependent kinases by p21. Mol Biol Cell 6: 387–400

Herbig U, Jobling W A, Chen B P C, Chen D J & Sedivy J M (2004) Telomere shortening triggers senescence of human cells through a pathway involving ATM, p53, and p21(CIP1), but not p16(INK4a). Mol Cell 14: 501–513

Houben J M J et al. (2009) Telomere shortening in chronic obstructive pulmonary disease. Respir Med 103: 230–236

Houssaini A et al (2018) mTOR pathway activation drives lung cell senescence and emphysema. J C I Insight 3

Hübner R H et al. (2008) Standardized quantification of pulmonary fibrosis in histological samples. BioTechniques 44: 507–511, 514–517

Kalucka J et al (2020) Single-Cell Transcriptome Atlas of Murine Endothelial Cells. Cell 180:764–779 e20

Kasahara Y. et al. (2001) Endothelial cell death and decreased expression of vascular endothelial growth factor and vascular endothelial growth factor receptor 2 in emphysema. Am J Respir Crit Care Med 163: 737–744

Kasahara Y et al (2000) Inhibition of VEGF receptors causes lung cell apoptosis and emphysema. J Clin Invest 106: 1311–1319

Kim N W & Wu F (1997) Advances in quantification and characterization of telomerase activity by the telomeric repeat amplification protocol (TRAP). Nucleic Acids Res 25: 2595–2597

Kim N W et al. (1994) Specific association of human telomerase activity with immortal cells and cancer. Science 266: 2011–2015

Kippin T E, Martens D J, van der Kooy D (2005) p21 loss compromises the relative quiescence of forebrain stem cell proliferation leading to exhaustion of their proliferation capacity. Genes Dev 19(6):756–767

Korsunsky I. et al. (2022) Cross-tissue, single-cell stromal atlas identifies shared pathological fibroblast phenotypes in four chronic inflammatory diseases. Med (N Y) 8: 3(7): 481–518 e14

Lai T P et al. (2017) A method for measuring the distribution of the shortest telomeres in cells and tissues. Nat Commun 8: 1356

Lingner J et al (1997) Reverse transcriptase motifs in the catalytic subunit of telomerase. Science 276: 561–567

Matmati S, Lambert S, Géli V & Coulon S (2020) Telomerase Repairs Collapsed Replication Forks at Telomeres. Cell Rep 30: 3312-3322.e3

Montandon M et al (2022) Telomerase is required for glomerular renewal in kidneys of adult mice. NPJ Regen Med 7(1): 15

Martínez P & Blasco M A (2017) Telomere-driven diseases and telomere-targeting therapies. J Cell Biol 216: 875–887

Niethamer T K, Stabler C T, Leach J P, Zepp J A, Morley M P, Babu A, Zhou S, Morrisey E E (2020) Defining the role of pulmonary endothelial cell heterogeneity in the response to acute lung injury. eLife 24: 9:e53072

Parrinello S et al (2003) Oxygen sensitivity severely limits the replicative lifespan of murine fibroblasts. Nat Cell Biol 5(8):741–7

Pettitt S J et al. (2009) Agouti C57BL/6N embryonic stem cells for mouse genetic resources. Nat Methods 6: 493–495

Piñeiro-Hermida S et al (2020) Telomerase treatment prevents lung profibrotic pathologies associated with physiological aging. J Cell Biol 219

Povedano J M et al. (2018) Therapeutic effects of telomerase in mice with pulmonary fibrosis induced by damage to the lungs and short telomeres. eLife 7

Povedano J M, Martinez P, Flores J M, Mulero F & Blasco M A (2015) Mice with Pulmonary Fibrosis Driven by Telomere Dysfunction. Cell Rep 12: 286–299

Pradad et al (2020) Promise of autologous CD34+ stem/progenitor cell therapy for treatment of cardiovascular disease. Cardiovascular Research 16:1424–1433.

Rafii S et al (2021) Reversal of emphysema by restoration of pulmonary endothelial cells. J Exp Med 2: 218(8): e20200938

Sarin KY et al (2005) Conditional telomerase induction causes proliferation of hair follicle stem cells. Nature 436(7053):1048–1052

Savale L et al (2009) Shortened telomeres in circulating leukocytes of patients with chronic obstructive pulmonary disease. Am J Respir Crit Care Med 179: 566–571

Stanley A J, Hasan I, Crockett A J, van Schayck O C P & Zwar N A (2014) COPD Diagnostic Questionnaire (CDQ) for selecting at-risk patients for spirometry: a cross-sectional study in Australian general practice. NPJ Prim Care Respir Med 24: 14024

Sharma G & Goodwin J (2006) Effect of aging on respiratory system physiology and immunology. Clin Interv Aging 1: 253–260

Shkreli M et al (2011) Reversible cell-cycle entry in adult kidney podocytes through regulated control of telomerase and Wnt signaling. Nat Med 18(1):111–9

Sidney L E, Branch M J, Dunphy S E, Dua H S, Hopkinson A (2014) Concise review: evidence for CD34 as a common marker for diverse progenitors. Stem Cells 32 (6):1380–9

Smogorzewska A & de Lange T (2002) Different telomere damage signaling pathways in human and mouse cells. EMBO J 21: 4338–4348

Sun H, Zhou Y, Fei L, Chen H & Guo G (2019) scMCA: A Tool to Define Mouse Cell Types Based on Single-Cell Digital Expression. Methods Mol Biol Clifton NJ 1935: 91–96

Tinkum K L et al. (2011) Bioluminescence Imaging Captures the Expression and Dynamics of Endogenous p21 Promoter Activity in Living Mice and Intact Cells. Mol Cell Biol 31: 3759–3772

Tsuji T, Aoshiba K & Nagai A (2010) Alveolar cell senescence exacerbates pulmonary inflammation in patients with chronic obstructive pulmonary disease. Respir Int Rev Thorac Dis 80: 59–70

Wang X, Wang R, Jiang L, Xu Q, Guo X. (2022) Endothelial repair by stem and progenitor cells. J Mol Cell Cardiol 163: 133–146

